# Inferring efficiency of translation initiation and elongation from ribosome profiling

**DOI:** 10.1101/719302

**Authors:** Juraj Szavits-Nossan, Luca Ciandrini

## Abstract

One of the main goals of ribosome profiling is to quantify the rate of protein synthesis at the level of translation. Here, we develop a method for inferring translation elongation kinetics from ribosome profiling data using recent advances in the mathematical modelling of mRNA translation. Our method distinguishes between the elongation rate intrinsic to the ribosome’s stepping cycle and the actual elongation rate that takes into account ribosome interference. This distinction allows us to quantify the extent of ribosomal collisions along the transcript and identify individual codons where ribosomal collisions are likely. When examining ribosome profiling in yeast, we observe that translation initiation and elongation are close to their optima, and traffic is minimised at the beginning of the transcript to favour ribosome recruitment. However, we find many individual sites of congestion along the mRNAs where the probability of ribosome interference can reach 50%. Our work provides new measures of translation initiation and elongation efficiencies, emphasising the importance of rating these two stages of translation separately.

## 1 Introduction

Understanding the rationale behind codon usage bias and the role of synonymous codons in regulating protein synthesis are among the main open questions in molecular biology. Despite the fact that mRNA translation is a pivotal stage of gene expression, its sequence determinants are in fact still largely elusive^1^. Recent advances in sequencing, such as ribosome profiling^2^, have made it possible to probe translation dynamics at codon resolution, allowing for quantitative studies of translational efficiency.

Ribosome profiling (Ribo-seq or ribosome footprinting as it is often called), is an experimental technique delivering a snapshot of ribosome positions along all transcripts in the cell at a given condition. Its archetypal version has been developed at the end of the 1960s to study translation initiation^3, 4^, and has been extended in the 1980s to investigate the role of slow codons and ribosome pausing^5^. Recently, Ingolia *et al*.^2^ revamped this technique to exploit the Next Generation Sequencing, and since then it is considered to be the state-of-the-art technique for studying gene expression at the level of translation.

In short, the method consists in isolating mRNA fragments (called “reads”) covered by a ribosome engaged in translation (∼ 30 nucleotides), which are then sequenced and aligned in order to build histograms of ribosome occupancy at codon resolution on each transcript. This technique has provided an unprecedented view on translation leading to many new discoveries^6^. Examples include detecting novel translation initiation sites^7^, identifying actively translated open reading frames^8^, quantifying the extent of stop codon readthrough^9^ and elucidating the translation of long non-coding RNAs^10^.

Translational activity on a given transcript is typically assessed by the number of read counts per kilobase of transcript per million mapped reads of the sample (RPKM), which takes into account the length of the transcript and the size of the sample. The RPKM is proportional to the ribosome density, which in turn is assumed to be proportional to the rate of translation – the more ribosomes on a transcript, the more efficient is protein synthesis. However, a large body of work based on mathematical modelling of ribosome dynamics suggests that the protein synthesis rate is negatively affected by increased ribosome density due to ribosome collisions^11–13^. To which extent ribosome collisions can be found using ribosome profiling has been an active topic of research^14–18^

One of the goals of ribosome profiling is to understand how the elongation rate along the transcript depends on the choice of codons. Codon elongation rates are usually estimated assuming that the ribosome density at codon *i* is proportional to the ribosome’s dwell time on that particular codon ^15, 16, 19–24^; this assumption follows from the conservation of the ribosome current assuming no ribosome drop-off. Our estimate of the drop-off rate of ∼ 10^−3^ s^−1^ (obtained from the probability of premature termination estimated to ≈ 10^−4^ per codon^25, 26^ and the elongation rate of the order of magnitude of ≈ 10 codon/s) justifies the hypothesis. The inferred elongation rates are then checked against mRNA codon sequence features such as codon usage bias, tRNA availability and mRNA secondary structures.

If ribosome collisions are not rare, then the elongation rates proxied by the inverse ribosome densities do not depend only on the molecular details of the elongation cycle, but also on the extent of slowing down due to ribosome traffic. The crux of the matter is that it is difficult to distinguish from the ribosome density alone whether the ribosome spent long time on a particular codon because of the long decoding time or because it had to wait for the downstream ribosome to move away. This distinction between the *actual* elongation rates that account for ribosome traffic and the *intrinsic* ones in the absence of other ribosomes has been well accounted for in the standard model for mRNA translation known as the totally asymmetric simple exclusion process (TASEP), which considers ribosomes moving along the transcript in a stochastic manner^11^. Yet, very few of the existing studies use the TASEP to infer elongation rates from Ribo-seq; ones that do either do not infer the intrinsic rates^24^ or use time consuming stochastic simulations to fit the Ribo-seq data^16, 22^.

In this work we develop an efficient method for inferring both actual and intrinsic codon-specific elongation rates from the ribosome profiling data based on the mathematical solution of the TASEP that we recently developed^12, 27^. Using the TASEP, we argue that the ribosome density alone is not sufficient to estimate the *absolute* elongation rates from the ribosome profiling data. Instead, our method infers elongation rates of an mRNA *relative* to the initiation rate of that transcript. Moreover, we propose new measures of translation efficiency that quantify the amount of ribosome traffic around the START codon and along the transcript. We apply our method to several Ribo-seq datasets in *Saccharomyces cerevisiae* and show evidence of local queuing *in vivo*.

## 2 Materials and methods

### 2.1 Ribosome profiling data

We have analysed publicly available ribosome profiling data of *Saccharomyces cerevisiae* from Guydosh *et al*.^14^, Pop *et al*.^20^ and Weinberg *et al*.^23^: NCBI GEO accession numbers GSM1279568, GSM1557447 and GSE75897 respectively. We downloaded the HDF5 files from Riboviz (https://riboviz.org)^28^ and mapped to A-site positions according the following table^29^:

**Table 1.** A-site locations for various footprint sizes.

| Fragment size | Frame 0 | Frame 1 | Frame 2 |
| --- | --- | --- | --- |
| 27 | 15 | 15 | 18 |
| 28 | 15 | 15 | 18 |
| 29 | 15 | X | 18 |
| 30 | 15 | 18 | 18 |
| 31 | 15 | 18 | 18 |
| 32 | X | 18 | 18 |
| 33 | 18 | 18 | 18 |

After obtaining the A-site read density profiles, our method successfully optimised 345 of the total 346 genes from the Guydosh dataset for which the experimental ribosome density necessary for normalisation was known from MacKay *et al*.^30^ Analogously, the optimisation was successful for 1051 out 1053 genes of the Pop dataset and for all 1589 genes in the Weinberg dataset. For the omitted genes the normalisation was not possible because it resulted in ribosome density larger than 1.

### 2.2 Notations

In this section we summarise the notations used in the paper. The main symbols for densities, rates, and rates relative to initiation are given in Table 2. When the quantity is codon specific we use the suffix *i* = 2, …, *L* to identify the codon number (the first codon after the START codon is at *i* = 2, the STOP codon is at *i* = *L*). Brackets {*·*} indicate a set of values: for instance {*a*_*i*_} is the set of all the values *a*_*i*_ for *i* = 2, …, *L*.

**Table 2.** Summary of the symbols used and their meaning.

| Symbol | Meaning |
| --- | --- |
| $L$ | length of the mRNA (in codons, including START) |
| $\ell$ | length of the ribosome (in codons) |
| $\alpha$ | initiation rate [$s^{-1}$] |
| $k_i$ | elongation rate [$s^{-1}$] of codon $i$, with $i = 2, \dots, L-1$ |
| $k_L$ (or $\beta$) | termination rate [$s^{-1}$] |
| $\{k_i\}$ | speed profile (elongation) of a given transcript |
| $\kappa_i = k_i/\alpha$ | relative (to initiation) elongation rate of codon $i$, or *elongation-to-initiation ratio* |
| $\{\kappa_i\}$ | relative (to initiation) elongation profile |
| $r_i$ | experimental (normalised) density of codon $i$ |
| $\{r_i\}$ | experimental (normalised) density profile |
| $r = \sum_{i=2}^L r_i / (L-1)$ | mean density of a given gene |
| $\rho_i$ | theoretical (normalised) density of codon $i$ |
| $\rho_i^{\text{ILA}}$ | theoretical (normalised) density of codon $i$ in the initiation-limited approximation (ILA) |
| $\{\rho_i\}$ | theoretical (normalised) density profile |
| $\rho_i^{\text{sim}}$ | simulated (normalised) density of codon $i$ |
| $\{\rho_i^{\text{sim}}\}$ | simulated (normalised) density profile |

We emphasise that we use *normalised* densities, in units of ribosomes (A-sites) per codon. The total density is thus the averaged ribosome profile 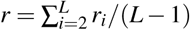, and the number of ribosomes translating an mRNA is *N* = *r*(*L*− 1).

### 2.3 Mathematical model for mRNA translation

We model translation by a stochastic process called the totally asymmetric simple exclusion process (TASEP) introduced by MacDonald *et al*.^11, 31^. The TASEP describes ribosome dynamics on a discrete one-dimensional lattice representing the coding part of the mRNA molecule. Each lattice site corresponds to a codon, and ribosomes cover *ℓ* = 10 sites, as the ribosome footprint covers ∼ 30 nt or equivalently ∼ 10 codons. Ribosomes on the lattice are tracked according to the position of their A-site. A codon *i* that is occupied by the A-site of the ribosome is labelled by *A*_*i*_ and is otherwise labelled by / ∅_*i*_.

A ribosome initiates translation at rate *α*, whereby its A-site is positioned at codon 2; this happens only if the codons 2, …, *ℓ* + 1 are not occupied by another ribosome’s A-site. The ribosome then advances from codon *i* to codon *i* + 1 at rate *k*_*i*_, provided that codon *i* + *ℓ* is not covered by the downstream ribosome (see top right drawing of the model in Fig. 1). We refer to *k*_*i*_ as the intrinsic elongation rate at which the ribosome advances in the absence of other ribosomes. Eventually, when the A-site of the ribosome is at the STOP codon (the *L*-th site), the ribosome detaches the mRNA at rate {*k*_*L*_ = *β*}. Each transcript in the model is then characterised by a set of *L* rates: initiation rate *α*, and elongation and termination rates {*k*_*i*_} = {*k*_2_, …, *k*_*L*_}.

**Figure 1.**
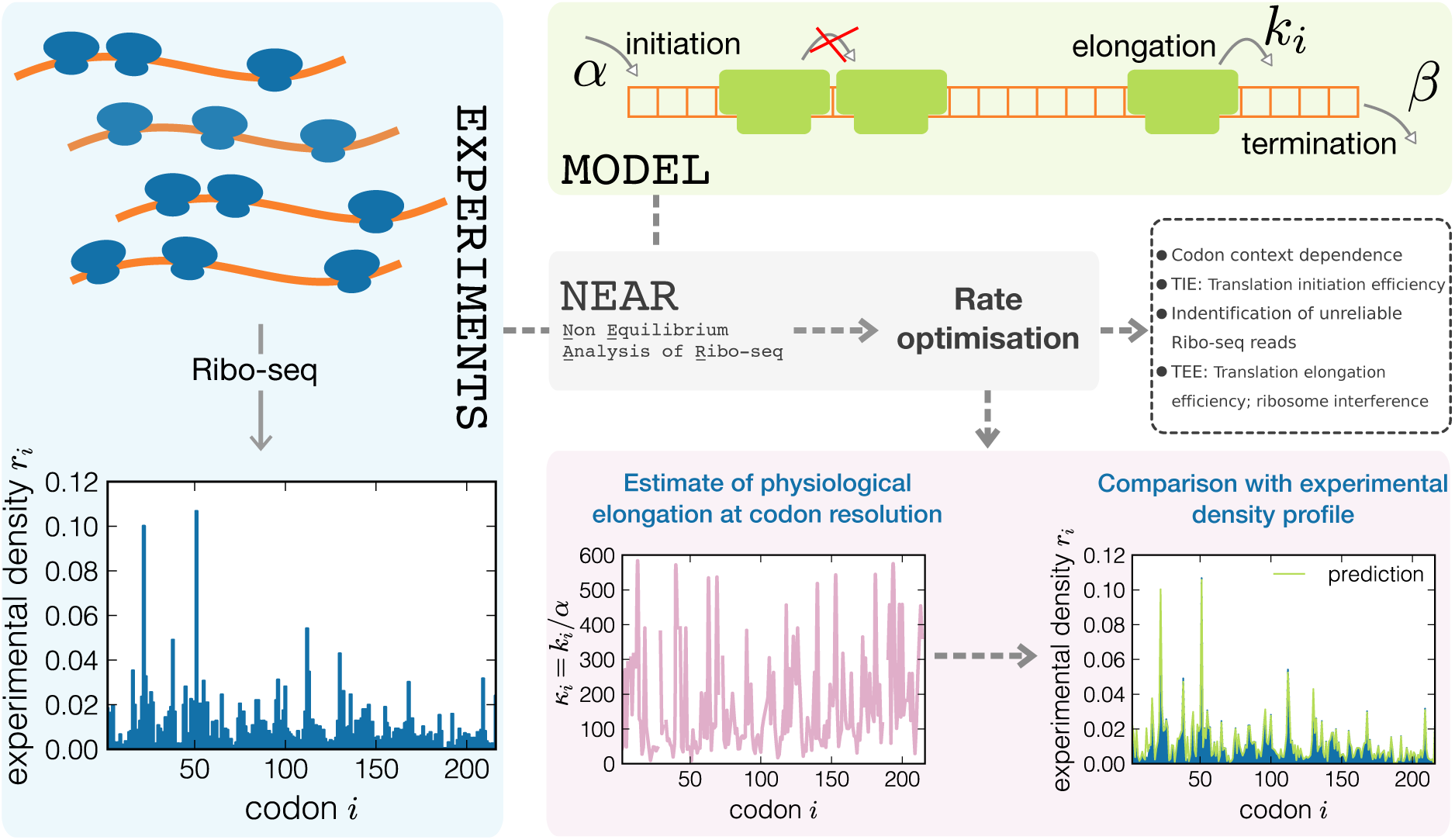
Sketch of the NEAR workflow for an individual gene (YAL007C). Experimental Ribo-seq profiles are first normalised and then analysed using the stochastic model. The normalised ribosome density profile {*r*_*i*_} is represented in the bottom left panel. The model is shown in the top right green box: ribosomes covering *ℓ* sites are added to the lattice with an initiation rate *α*, provided that the first *ℓ* sites are not occupied by the A-site of another ribosome. Ribosomes then move from site *i* to site *i* + 1 at rate *k*_*i*_ provided that the A-site of the neighbouring ribosome downstream is at least *ℓ* sites away. Eventually, ribosomes leave the lattice at rate *β* (*k*_*L*_ = *β*) when their A-site is on the last site. In this drawing *ℓ* = 3 for clarity, while in our analysis we used *ℓ* = 10. NEAR searches for the optimal elongation rates *k*_*i*_ (relative to initiation rate *α*) for which the stochastic model reproduces the experimental ribosome profile. Once we find the optimal rates, we examine the extent of ribosome traffic using the translation initiation and elongation efficiencies (TIE and TEE), analyse the context dependence of elongation rates, and identify problematic transcript regions in which Ribo-seq data are not consistent with the model.

The process is described by the probability density *P*(*C, t*) to find a configuration *C* of ribosomes on an mRNA at a particular time *t*. By *configuration* we mean a particular arrangement of ribosomes described by the positions {*A*_*i*_} of their A-sites. The time evolution of *P*(*C, t*) is governed by the master equation

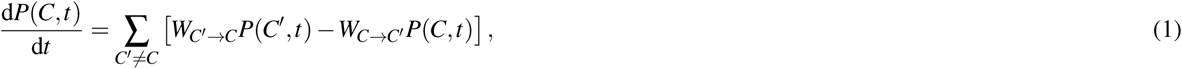

where *W*_*C*→*C*′_ is the rate of transition from *C* to *C*′. We assume that translation takes place in the stationary limit in which case Eq. (1) becomes a system of linear equations,

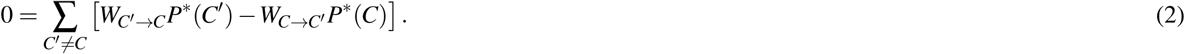

The three main quantities of interest are the rate of translation *J*, which measures the amount of proteins produced per unit time, the local ribosome densities *ρ*_*i*_, which measure the probability of detecting a ribosome at codon *i*, and total ribosome density *ρ*, which measures the average number of ribosomes per unit length of the transcript (in codons). In the stationary TASEP, *J, ρ*_*i*_ and *ρ* are defined as

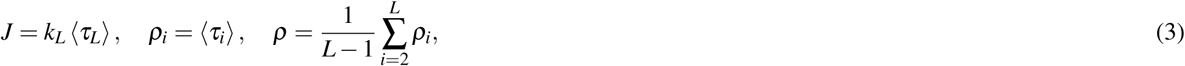

where averaging is performed with respect to the steady-state probability *P*^∗^(*C*) and *τ*_*i*_ is an occupation number whose value is equal to 1 if codon *i* is occupied by the A-site of the ribosome and is 0 otherwise. If we ignore premature termination due to ribosome drop-off, then *J* is constant across the transcript and is equal to the actual rate at which ribosomes initiate translation,

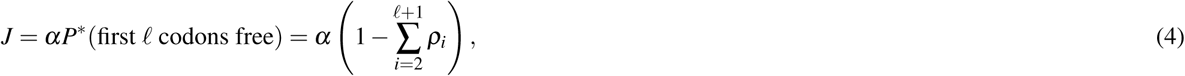

where *P*^∗^(first *ℓ* codons free) is the steady-state probability that codons 2, …, *ℓ* + 1 are not occupied by an A-site.

Computing these quantities requires an exact knowledge of *P*^∗^(*C*), which is known only in the biologically unrealistic case of *ℓ* = 1 and uniform elongation rates^32^. Instead, we compute *J, ρ*_*i*_ and *ρ* using two approximation methods: the mean-field approximation developed in MacDonald *et al*.^11, 31^ and initiation-limited approximation developed in Szavits-Nossan *et al*^12, 27^. Details of these methods are presented in Supplementary Information.

### 2.4 Computer program

Computer program (NEAR) for inferring elongation rates from ribosome profiling data is available under GNU General Public License v3.0 at https://github.com/jszavits/NEAR.

## 3 Results

We base our method on a well-established stochastic model for mRNA translation, the *totally asymmetric simple exclusion process* (TASEP), which we describe in detail in Materials and Methods. Over the years, the model has been improved in many ways to better match real translation^33, 34^ and has been repeatedly used to interpret experimental data^16, 18, 24, 35–38^.

In principle, the knowledge of initiation, elongation and termination rates allows one to compute simulated ribosome density profiles and protein production rates that can be compared to experimental outcomes. However, there is an open debate regarding the estimates of these rates, and no direct experimental method to measure them exists. For example, codon-specific translation elongation rates *k*_*i*_ are often assumed to be proportional to the tRNA gene copy number (GCN) or to the local tRNA adaptation index (tAI) ^39–41^.

Here we take a different approach and use the model to quantitatively determine codon elongation rates from ribosome profiling data. This is an *inverse problem*, since we need to optimise the inputs (parameters *α* and {*k*_*i*_}) in order to match the outputs (Ribo-seq data). There are three main difficulties in solving this problem, which we discuss below.

1. The parameter space is extremely vast. A typical protein consists of a several hundreds of amino acids, meaning that one generally needs to optimise a comparable number of parameters.
2. There is a complex non-linear relation between the set of rates {*k*_*i*_} and the ribosome density profile. A change in a single *k*_*i*_ may affect a large part of the density profile.
3. Ribosome density profile predicted by the stochastic model depends only on the ratios between the elongation rates and the initiation rate, meaning that it is not possible to estimate *absolute* rates without integrating more information.

We now explain how our method tackles these problems and how it compares to existing methods that have been proposed to infer ribosome dynamics from ribosome profiling data^16, 20, 22, 24^.

Our method searches for optimal elongation rates at each codon position and separately for every transcript, i.e. we do not reduce the parameter space by assuming equal elongation rates for every instance of the same codon^20, 22^. Importantly, we use an analytic expression for the ribosome density profile that we recently derived in the initiation-limited regime^12, 27^. This relationship allows for a quick computation of the ribosome density profile instead of running costly stochastic simulations for every iteration of the optimisation process^16, 22^. Furthermore, we emphasise that our method infers *intrinsic* elongation rates (relative to the initiation rate) related to the ribosomal elongation cycle, which may differ from the *actual* elongation rates that also take into account slowing down due to ribosome traffic^24^. Thus, we are able to detect separately the mean decoding time for a particular codon and the mean time that the ribosome spends waiting for a ribosome downstream of it to move away. This distinction is central to our method.

Before we present further details of our method, we first discuss the problem of estimating absolute elongation rates, which limits the amount of information that can be inferred from ribosome profiling data alone.

### 3.1 Ribosome profiles alone cannot estimate absolute elongation rates

We remind that the ribosome density *ρ*_*i*_ measures the probability of detecting a ribosome’s A-site at codon position *i* (see Methods and Materials for further details). In the Supplementary Information we show that *ρ*_*i*_ depends only on the ratios between the elongation/termination rates {*k* _*j*_} and the initiation rate *α*–we will refer to these ratios as {*κ*_*j*_}. Thus given the ribosome densities {*ρ*_*j*_}, one can only infer the ratios {*κ*_*j*_}, but not the absolute rates {*k* _*j*_} and *α*. Since the initiation rate *α* is highly gene-dependent, it is not possible to compare the elongation-to-initiation ratios {*κ*_*j*_} from different genes without the knowledge of *α* for each gene. We demonstrate this point in Fig. 2(a), which shows the outcome of simulations of translation having different absolute rates {*k*_*i*_} but same relative speed profile {*κ*_*i*_}.

**Figure 2.**
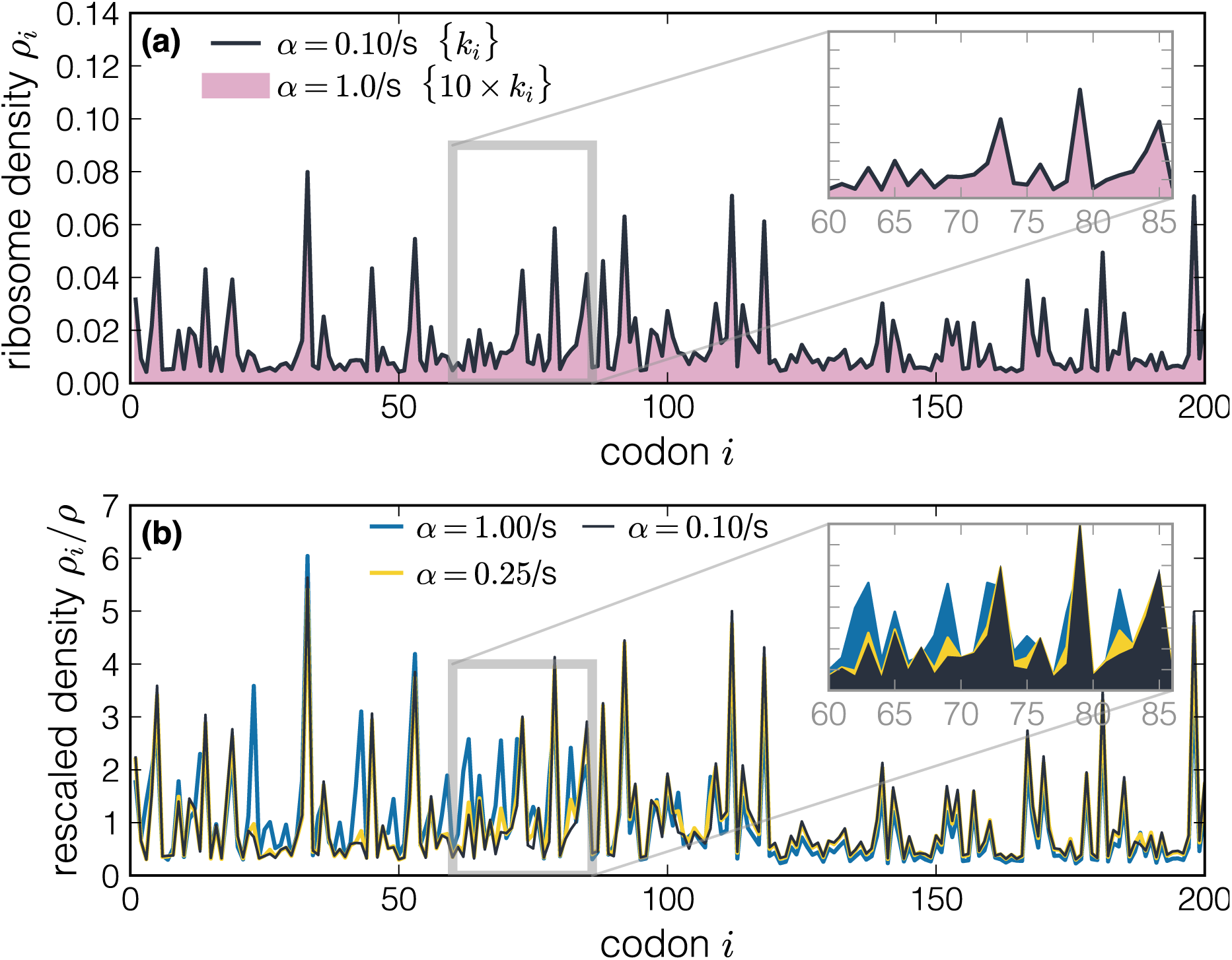
*in silico* density profiles. In panel (a), black line, we show the density profile obtained from the stochastic simulation of a transcript with a speed profile {*k*_*i*_} and initiation rate *α* = 0.1/s. The pink region corresponds to the profile of a transcript simulated with a tenfold larger initiation rate, but keeping {*κ*_*i*_} constant (i.e. also increasing the elongation rates by a factor 10). This shows that densities obtained with the same elongation-to-initiation ratios {*κ*_*i*_} are indistinguishable. In panel (b) we fix the speed profile {*k*_*i*_} for three different values of the initiation rate *α* and we plot the rescaled profiles *ρ*_*i*_*/ρ*. As expected, by increasing the initiation rate we obtain different profiles with increasing density and traffic effects.

We now examine two approaches that have been proposed to deal with this problem. The first approach is to fix a unique timescale shared by all mRNAs, for instance the average ribosome speed^42^ or the average codon decoding time^24^, which in turn allows one to estimate the initiation rate for each gene. The second approach is to normalise *ρ*_*i*_ by the average ribosome density *ρ* for that gene. This is a common practice in the analysis of Ribo-seq data, whereby the ribosome footprint read densities on individual codons are normalised by the average ribosome footprint density for that gene. The scaled read density is then assumed to be independent of the initiation rate, allowing for different genes to be compared. We argue that both of these approaches are problematic. In the first approach, the average elongation rate or the average decoding time could be highly variable from transcript to transcript, which in turn would introduce a bias when comparing absolute elongation rates between different genes. In the second approach, the normalisation of *ρ*_*i*_ by *ρ* does not necessarily mean that genes with different initiation rates can be directly compared. We show that explicitly by computing *ρ*_*i*_*/ρ* in our model for different initiation rates but keeping the elongation speed profile {*k*_*i*_} fixed. As shown in Fig. 2(b), we find qualitatively different profiles for different initiation rates. This observation is further supported by the analytic expression for *ρ*_*i*_, which predicts a non-linear *α*-dependent correction to the linear expression *ρ*_*i*_ ≈ *α/k*_*i*_ (see Supplementary Information).

Instead, our approach is to scale *κ*_*i*_ by the termination-to-initiation ratio *κ*_*L*_ which removes dependence on the initiation rate since *κ*_*i*_*/κ*_*L*_ = *k*_*i*_*/k*_*L*_. Later we show that the values of *κ*_*L*_ inferred from ribosome profiling data in *S. cerevisiae* have among the least variation of all codons, which supports our choice for *κ*_*L*_ as the scaling factor. In addition, we introduce new measures of translation efficiency and ribosome traffic that take values between 0 and 1 and can be compared between different genes.

### 3.2 Non-Equilibrium Analysis of Ribo-seq (NEAR)

After we have shown that the ribosome density profile alone can inform us only on the ratios {*κ*_*i*_} between the elongation rates and the initiation rate, we now turn to the method for inferring {*κ*_*i*_} from Ribo-seq. We call the method Non-Equilibrium Analysis of Ribo-seq data (NEAR) because the model (the TASEP) that we use is borrowed from nonequilibrium statistical mechanics.

NEAR infers {*κ*_*i*_} with an optimisation procedure that aims to find a model-predidcted density profile *ρ*_*i*_ which is a close match to the experimental one {*r*_*i*_} (see Fig. 1). This is possible since we have recently found a mathematical expression for the ribosome density profile in terms of translation initiation, elongation and termination rates. This expression was obtained under the assumption of a limiting initiation rate *α*^12, 27^, which is supposed to hold for most of the mRNAs under physiological conditions (see Supplementary Information). However, we emphasise that the initiation-limited approximation does not assume that ribosome collisions are absent. Instead, our analytic solution takes ribosome collisions into account and is applicable to a wide range of initiation rates as long as they are smaller than the elongation and termination rates (see Supplementary Information).

We have applied our method to ribosome profiling data of *Saccharomyces cerevisiae* obtained by Guydosh *et al*.^14^, Pop *et al*.^20^ and Weinberg *et al*.^23^. These datasets were selected for their lack of using cycloheximide to inhibit translation elongation, which is known to distort ribosome coverage profiles^43, 44^. The raw data was processed by the Riboviz software^28^ and mapped to A-site positions following the table provided in Ahmed *et al*.^29^. After obtaining the A-site read density profiles the method proceeded in four steps, which we summarise below.

1. We first normalised the number of A-site reads on each codon by the total number of reads mapped to the transcript. This number was then multiplied by the absolute ribosome density for that particular gene obtained by polysome profiling experiments in MacKay *et al*.^30^. The end result is a normalised ribosome density profile {*r*_*i*_} that reveals how likely is to find a ribosome at codon *i*.
2. Next, we solved a least-squares optimisation problem which consisted of finding {*κ*_*i*_} that minimise the objective function

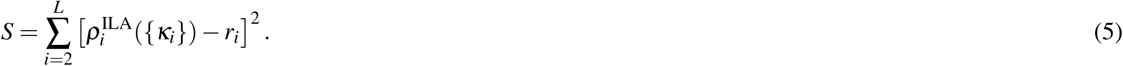

Here 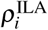 is the model-predicted ribosome density in the initiation-limited approximation (ILA). The starting point for optimisation were {*κ*_*i*_} obtained from the mean-field solution of the exclusion process. Details of 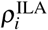 and the mean-field solution are presented in Supplementary Information.
3. Once we found the best estimate of {*κ*_*i*_}, we then computed the exact density profile from stochastic simulations using the estimated {*κ*_*i*_}. We note that the simulated density may be different from the analytic density if the initiation rate is too high, which we checked in the next step.
4. In the last step we performed two quality checks on each *κ*_*i*_ obtained by least-squares optimisation:
  a. We first verified that the initiation-limited approximation was applicable by comparing the analytic density with the simulated one. This step is necessary because our solution of the model is approximate and may not be valid if the initiation rate is too high, see Refs.^12, 27^ and also the Supplementary Information. We accepted *κ*_*i*_ if the relative error between the analytic and simulated density was smaller than 10 %. If not, we repeated the check using the value of *κ*_*i*_ obtained in the mean-field approximation.
  b. For those *κ*_*i*_ that passed the previous check, we verified that the simulated density reproduced the experimental density *r*_*i*_ (within 5% tolerance).

These are the main steps of NEAR, and we provide further mathematical details in the Supplementary Information.

We emphasise the importance of optimising the absolute ribosome densities {*ρ*_*i*_} (step 2), rather than the scaled ones, {*ρ*_*i*_*/ρ*}, as in other methods that analyse ribosome profiling data^16, 20^. The problem is that {*ρ*_*i*_*/ρ*} remains the same if we multiply all *ρ*_*i*_ by a constant factor, which in turn means that different density profiles {*ρ*_*i*_} can result in the same scaled profile {*ρ*_*i*_*/ρ*}. Since each *ρ*_*i*_ is uniquely determined by the set of elongation-to-initiation ratios {*κ*_*i*_}, we conclude that the scaled density profile {*ρ*_*i*_*/ρ*} does not uniquely determine {*κ*_*i*_}, see for instance Figure S2. Thus, we reiterate that ribosome profiling data is not sufficient to infer ribosome dynamics and in turn the extent of ribosome traffic without the additional information on the mean number of ribosomes bound per mRNA (step 1).

Our quality check in step 4 is also able to reject codons whose *κ*_*i*_ cannot be trusted, and identify why the inference of elongation rates for those codons is problematic. Importantly, we are able to distinguish whether our analytic solution is satisfied or not (point 4(a)), or if the problem is due to the model being inconsistent with the experimental data (point 4(b)).

Before moving on to real sequences of *S. cerevisiae*, we tested NEAR on a mock sequence with known (*k*_*i*_} (Figure S3(a)), and checked that it can accurately infer the original elongation rates provided the initiation rate is not too high (Figure S3(b) and (c)). We also remark that the quality check allows us to push the analysis to relatively high initiation rates (Figure S3(d)). In those cases, however, the number of rejected codons may become significant. For a very high initiation rate we expect the initiation-limited approximation to fail in which case NEAR resorts to the mean-field approximation, whose estimates are further verified.

#### 3.2.1 Using NEAR to study translation of individual genes

We demonstrate our method on a particular gene (YLR301W) using Ribo-seq data from the Weinberg dataset^23^. We first compute the normalised experimental density profile {*r*_*i*_} using the experimental absolute density *r* from MacKay *et al*.^30^ (in units of ribosomes/codon). This profile is then analysed following the method explained in the previous section. A set of elongation-to-initiation ratios {*κ*_*i*_} is obtained by optimising the match between the model-predicted density profile and the experimental one. Each *κ*_*i*_ is then examined to see whether it provides a good prediction for that particular codon position and to check for inconsistencies in the method as previously explained. There are few values of *κ*_*i*_ that do not pass this quality check, which are rejected and are not included in the final analysis. This is a typical example, though for some genes the fraction of rejected codons is substantial and our inference procedure may be less reliable. We will come back to this point later.

In Fig. 3(a) we plot the optimised {*κ*_*i*_} profile that passed the quality check (blue line, triangle markers) compared to the naive estimate 1*/r*_*i*_ (orange line, round marker) that ignores ribosome interference but it is usually judged as a good estimator of the elongation rate. We find many codon positions where the two profiles {*κ*_*i*_} and {1*/r*_*i*_} significantly differ from each other. Moreover, we identify values of *κ*_*i*_ that are not consistent with the model, while this cannot be done when using {1*/r*_*i*_} as a proxy for elongation determinants.

**Figure 3.**
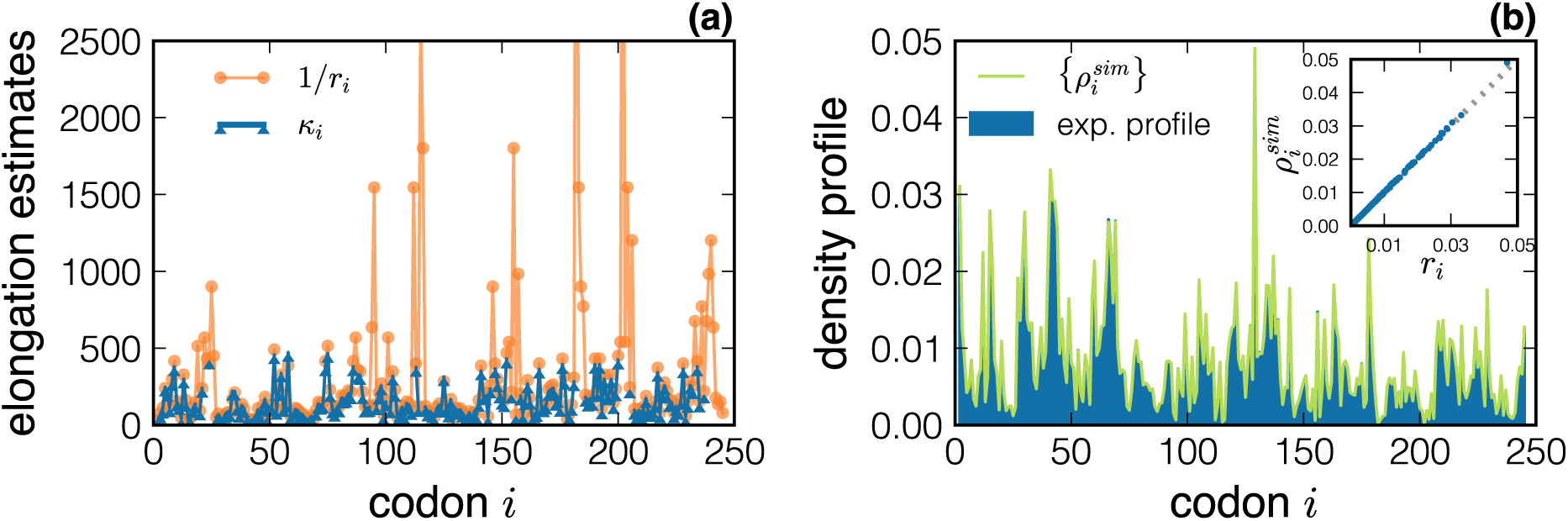
Results of NEAR applied to the YLR301W gene. (a) The optimised profile {*κ*_*i*_} is plotted (blue line, triangle markers) as a function of the codon position *i*, and compared to the naive estimates {1*/r*_*i*_} (orange line, round marker). In panel (b) we compare the model-predicted density profile obtained using the inferred {*κ*_*i*_} (green line) with the experimental normalised profile {*r*_*i*_}. The inset shows the scatter plot between the two densities (for each codon *i*) demonstrating an excellent agreement between theory and experiments.

The result of a stochastic simulation of ribosome dynamics performed with the optimised elongation ratios {*κ*_*i*_} is shown in Fig. 3(b). The agreement between the simulated density profile 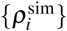 (green line) and the experimental one (in blue) is excellent for most of the codons. The inset shows the scatter plot between the values (for each codon) of the simulated and experimental ribosome density.

### 3.3 Estimate of elongation-to-initiation ratios at codon resolution in yeast

We analysed three different datasets^14, 20, 23^ and gathered the NEAR elongation-to-initiation ratios {*κ*_*i*_} for each gene. The percentage of codons that passed the quality check (points 4(a) and (b)) for the Weinberg, Pop and Guydosh datasets is 75%, 66% and 44%, respectively. These are the percentages of the total number of analysed codons, i.e. without taking into account different transcript lengths.

We also computed the percentage of rejected codons for each transcript. The percentages of codons that were rejected at point 4(a) have a median value of 2.3% (Weinberg), 3.8% (Pop) and 8.2% (Guydosh). The respective medians for the percentages of codons that passed 4(a) but were rejected at point 4(b) are 9.5%, 16% and 26.7%. Again, the best fit is achieved for the Weinberg dataset.

We note that our analysis included only transcripts with large number of reads per codon (10 or more), i.e. with high ribosome traffic. If we had analysed all transcripts, the percentage of accepted codons would have been higher. However, many transcripts with low read count have codons with zero reads, which are difficult to handle in the model as they imply unphysically large elongation speed.

We now turn to the codons that passed the quality check. The estimated elongation-to-initiation ratios passing the quality check are plotted in Fig. 4 and compared to the naive estimates 1*/r*_*i*_. In particular, we find many instances where 1*/r*_*i*_ deviates from *κ*_*i*_ obtained by NEAR. The model predicts that *κ*_*i*_ ≈ 1*/r*_*i*_ if there are very few ribosomes on the transcript so that ribosome collisions are rare. Our findings in Fig. 4 thus suggest that the effect of ribosome interference is not negligible. We will discuss this point later when we introduce better measures for detecting ribosome interference.

**Figure 4.**
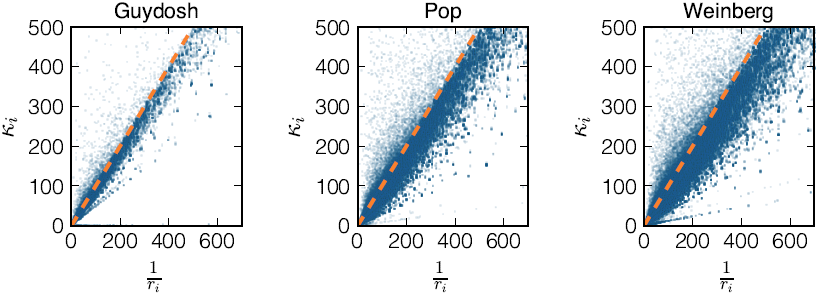
Scatter plot of the elongation-to-initiation *κ*_*i*_ for each codon that passed the quality check versus the inverse of the experimental density 1*/r*_*i*_ for the Guydosh, Pop and Weinberg datasets. The dashed line corresponds to the bisect.

Next, we wanted to understand if each codon type has a characteristic decoding time and verify or reject a common hypothesis that elongation rates are determined by the availability of aminoacyl-tRNAs. By definition, *κ*_*i*_ is equal to the ratio *k*_*i*_*/α* between the elongation rate of codon *i* and the initiation rate *α* of the gene. Because the initiation rates are likely to be gene-specific, we do not know if the observed variation in elongation-to-initiation ratios of the same codon types (see Figure S4) is due to variation in the elongation or initiation rates.

However, we observe that STOP codons show the least variability of all the elongation-to-initiation ratios {*κ*_*i*_} in the Guydosh and Weinberg datasets (Figure S4). This result is consistent with the expectation of a context-independent termination. Thanks to this observation, we then compute the elongation-to-termination ratio *κ*_*i*_*/κ*_*L*_ = *k*_*i*_*/β*, i.e. the elongation rate of codon *i* with respect to the termination of the gene under investigation (Figure S5). This quantity does not depend on the initiation rate *α* that is likely to be context-dependent and different from gene to gene. Indeed, the variation in *κ*_*i*_*/κ*_*L*_ linked to the same codon type is now more uniform across 61 codon types, especially in the Guydosh dataset (Fig. S5). We have also compared median values of *κ*_*i*_*/κ*_*L*_ for each codon type against two common measures of tRNA availability: a codon-dependent rate of translation based on the tRNA gene copy number (GCN) corrected for the wobble base pairing from Weinberg *et al*.^23^, and the tRNA adaptation index (tAI)^45^. We find a moderate correlation between the median of the *κ*_*i*_*/κ*_*L*_ distributions and the corresponding tRNA gene copy numbers (Figure S6). This result suggests that the elongation speed of individual codons is only partially determined by their codon type.

We now turn to ribosome traffic and its impact on translation efficiency. In the following sections we will define quantities that, contrary to the *κ*_*i*_, can be used to compare translation efficiency of different genes. Those quantities, which we name the translation initiation efficiency (TIE) and the translation elongation efficiency (TEE), can be used to rank initiation of different transcripts and quantify the impact of ribosome interference along a mRNA.

### 3.4 Translation Initiation Efficiency (TIE)

By running stochastic simulations with the inferred *κ*_*i*_ we can measure the ribosomal current *J* divided by the initiation rate *α*, which is a quantity dependent on {*κ*_*i*_} only; the current *J* can be used as a proxy for protein synthesis rate per mRNA.

In the biological literature translation initiation is often identified with protein synthesis rate, i.e. *J* = *α*. However this is true only if initiation is much slower than elongation so that essentially only one ribosome is translating a transcript at a given time. Yet, this approximation is too crude to quantitatively describe translation^12^. Instead, when more than one ribosome is engaged in translation, *J* becomes a function of *α* and the elongation rates {*k*_*i*_}; the current *J* can be thought of as the intrinsic initiation rate *α* multiplied by the probability that the first codons of the mRNA are not occupied by another ribosome (which would otherwise obstruct initiation).

Therefore we propose to use *J/α* as a measure of the Translation Initiation Efficiency (TIE), which takes values between 0 and 1. The TIE would be equal to 1 in the optimal case in which initiation is not hindered by ribosome traffic (*J* = *α*, hence TIE = 1). Otherwise, the TIE gives the probability that the first codons, potentially interfering with ribosome recruitment and initiation, are unoccupied. A TIE smaller than 0.5 means that more than half of the times a new ribosome tries to initiate, it fails because of another ribosome whose A-site is located within the first ten codons. In Fig. 5 we plot the histograms of TIE for all the genes and datasets included in our study. We find that almost all genes show TIE > 0.5 with a median value around 0.8 for all the datasets. These values suggest that the first codons are mainly free from ribosomes that are already engaged in translation.

**Figure 5.**
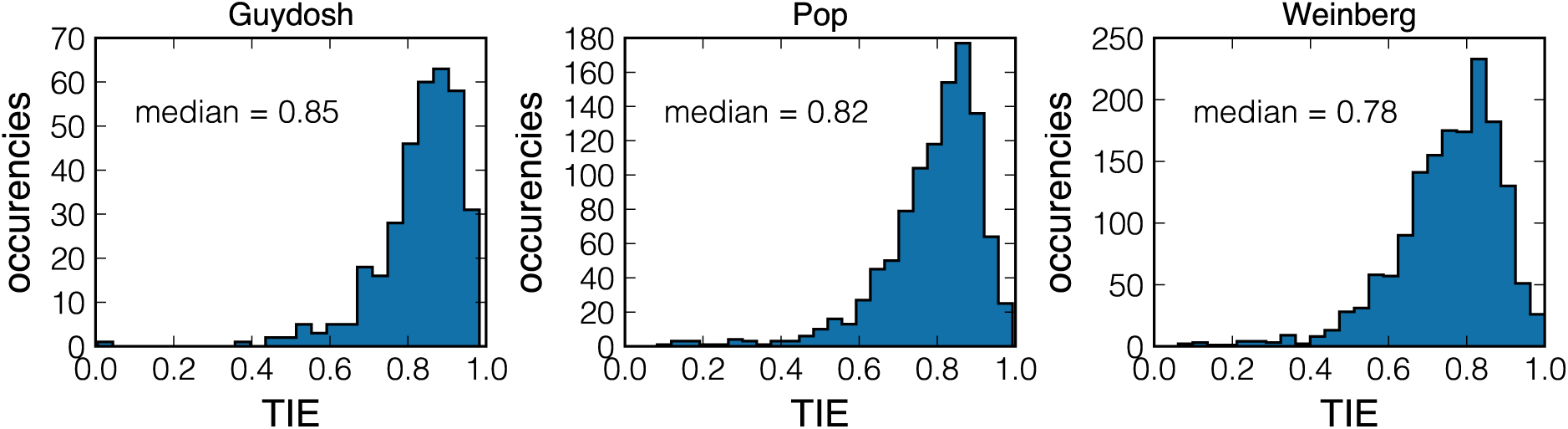
Histogram of estimated Translation Initiation Efficiency (TIE) for all *Saccharomyces cerevisiae* genes included in our study for the Guydosh, Pop and Weinberg datasets. The TIE gives the probability that the first codons are unoccupied.

Our previous theoretical work on the exclusion process showed that if translation is rate-limited by initiation, then TIE predominately depends on the *κ*_*i*_ of the first *ℓ* ≈ 10 codons, which is the ribosome footprint on the mRNA^12^. Based on that prediction, TIE > 0.5 is a strong signature that the codon sequences, and in particular the first codons of *Saccharomyces cerevisiae* genes might have been selected to optimise translation initiation.

### 3.5 Translation Elongation Efficiency (TEE) shows congestion of ribosomes *in vivo*

In contrast to the TIE, we define an efficiency index for translation elongation that identifies local ribosome interference along the transcript and not only around the initiation region. In order to do that, we emphasise that the total time *t*_*i*_(total) that a ribosome spends with its A-site on a given codon *i* can be seen as the sum of two contributions: the time *t*_*i*_(intrinsic) needed to decode this codon and incorporate the new amino acid to the growing peptide chain, plus the time *t*_*i*_(collision) spent in a queue waiting for the downstream ribosome to move. For each codon *i, t*_*i*_(total) = *t*_*i*_(intrinsic) + *t*_*i*_(collision). The inverse of *t*_*i*_(total) is the actual elongation rate, while the inverse of *t*_*i*_(intrinsic) is the intrinsic elongation rate *k*_*i*_, i.e. the elongation rate in the absence of other ribosomes. The distinction between these two rates is important because the actual elongation rates may be much smaller than the intrinsic ones in genes with higher initiation rates in which ribosomal collisions are more likely. Thus analysing the actual instead of intrinsic elongation rates could obscure our search for the molecular determinants of the translation speed.

We consider a codon as *efficient* if a ribosome attempting to translate it is not blocked by other ribosomes. We thus define the Translation Elongation Efficiency at codon *i* (TEE_*i*_) as the ratio of intrinsic and total time: TEE_*i*_ = *t*_*i*_(intrinsic)*/t*_*i*_(total), or put differently, as the ratio between the actual and intrinsic elongation rate. The TEE_*i*_ is a measure of the local mRNA congestion seen by a ribosome translating the codon *i* and it depends on the context at which the codon is placed. Mathematically, it is equivalent to the probability that the *i* + 1 … *i* + *ℓ* codons are not occupied, given that a ribosome’s A-site is at site *i*. If the intrinsic decoding time of the ribosome is equal to the total time dwelt on the codon, then the ribosome experiences on average no interference with other ribosomes and TEE_*i*_ = 1. Otherwise, 0 *<* TEE *<* 1. In the extreme case of the completely jammed ribosome one would get TEE_*i*_ ≈ 0, i.e. the ribosome is ready to advance but it is not allowed to move forward because the transcript is overcrowded. Furthermore there is a relationship between the TEE_*i*_ and TIE given by TEE_*i*_ = TIE*/*(*κ*_*i*_*ρ*_*i*_). Further details are given in Supplementary Information.

We note that the TEE_*i*_ is a function of {*κ*_*i*_} only, meaning that ribosome interference is governed by the balance between initiation and elongation rates. A TEE profile that is close to 1 means that initiation is not frequent enough to cause ribosome congestion along the transcript. Inferring TEE profile from ribosome profiling data is thus a convenient method for testing whether translation is limited by initiation. We also stress that both the TIE and TEE_*i*_ are dimensionless quantities that take values between 0 and 1. Therefore it is possible to compare the TIE and TEE profiles between different genes.

In Figure 6(a)-(b)-(c)-(d) we plot the TEE profile of four randomly selected genes. We observe that the TEE is typically close to 1 indicating that traffic is negligible for most of codons. We also identify particular codons where ribosome interference is significant and TEE_*i*_ drops to 0.6. These examples demonstrate that NEAR can locate, at codon resolution and excluding unreliable estimates, particular regions on the transcript that are affected by ribosome interference.

**Figure 6.**
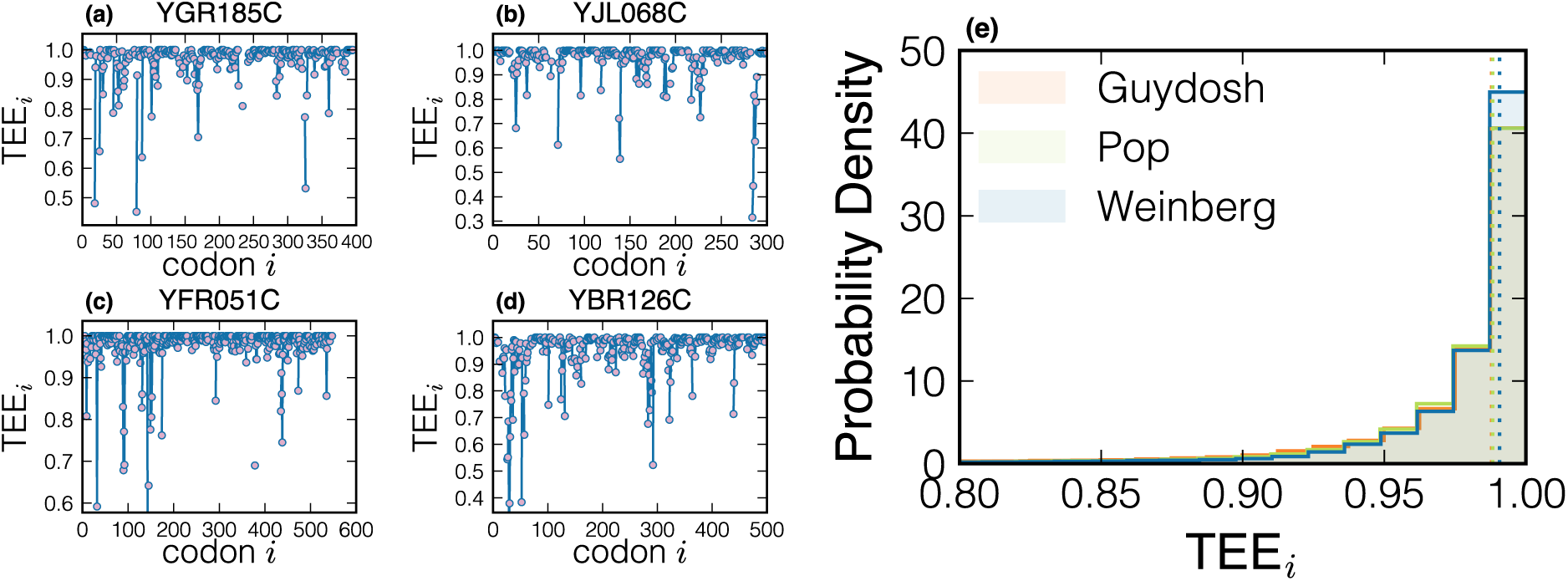
Panels (a)-(b)-(c)-(d) show single-gene TEE profiles (randomly selected genes from Weinberg dataset). Each point represents a codon, blue lines connects adjacent codons. Isolated points mean that their neighbouring codons have been rejected. In panel (e) we plot the distributions of the TEE collected on all codons of the three datasets analysed. The dashed vertical lines represent the median of those distributions (Guydosh: 0.988, Pop: 0.988, Weinberg: 0.990).

After analysing TEE profiles of all genes included in our study, we observe that the distribution of TEE_*i*_ for all the codons that passed the NEAR quality check is peaked at 1, with the median at about 0.99, as shown in Fig. 6(e). This result is consistently found in all three datasets that we analysed, suggesting that ribosome interference is present only locally on a few codons, and is generally absent *in vivo*.

If the ribosome density on a given transcript is high, one would expect to see an increased number of ribosomal collisions resulting in the TEE profile that clearly deviates from 1. In Figure 7 we present the mean of the TEE profile for each gene that we analysed compared to the ratio of the ribosome density for that gene and the maximum achievable density *ρ*_max_ = 1*/ℓ*, where *ℓ* ≈ 10 codons is the ribosome footprint length. The results across all three Ribo-seq datasets clearly show that genes with low ribosome density have the mean TEE very close to 1 (few collisions). On the other hand, the mean TEE of genes with high ribosome density deviates significantly from 1 (many collisions).

**Figure 7.**
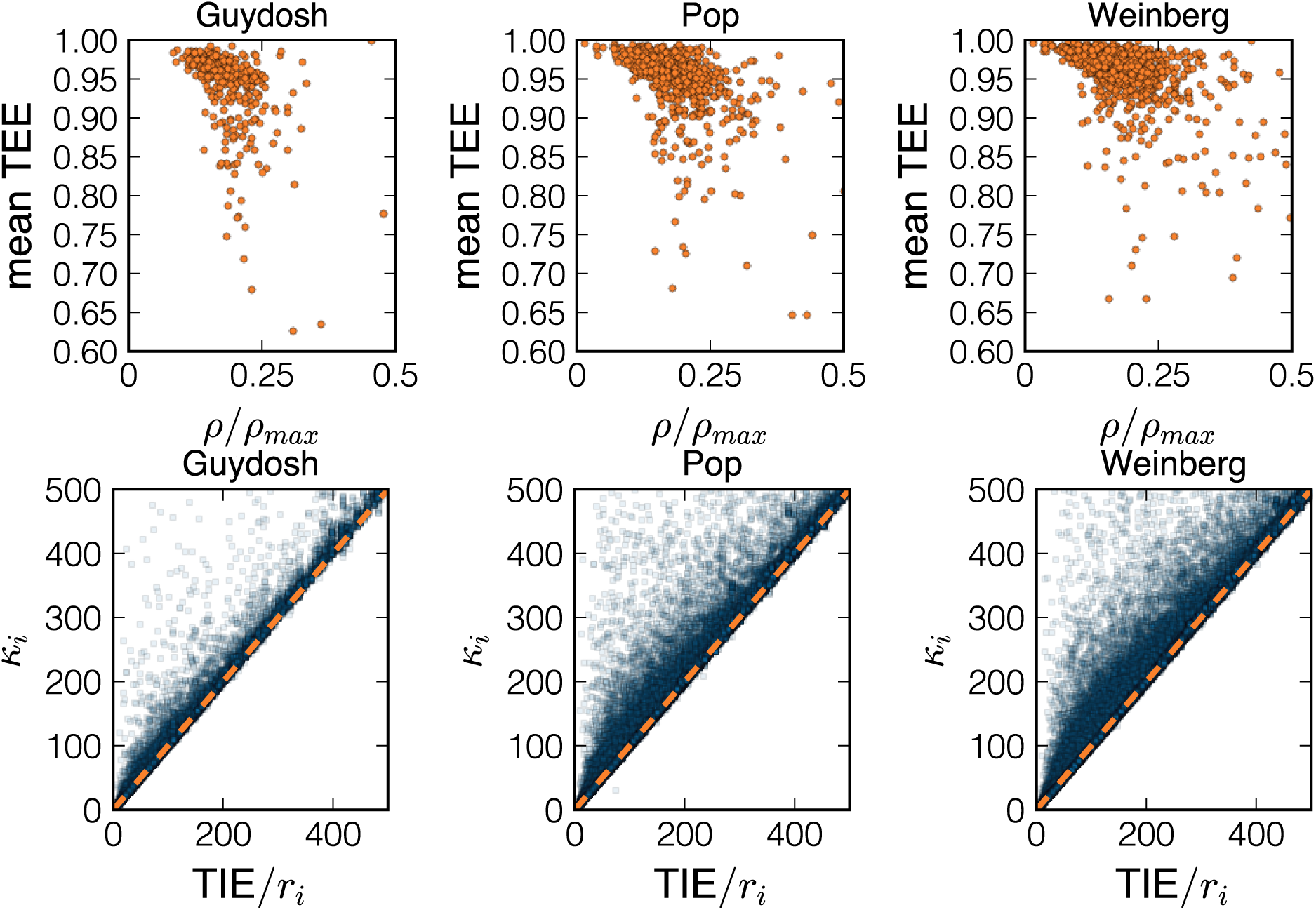
In the first row we show the ribosome density *ρ* normalised with the maximal density *ρ*_max_ versus the mean value of TEE along the transcript. In the second raw we plot *TIE/r*_*i*_ versus *κ*_*i*_ to emphasise the extent of traffic in determining the ribosome’s dwelling time. The orange dashed line is the bisect *TIE/r*_*i*_ = *κ*_*i*_ (no traffic). Codons far from that line are the ones more impacted by ribosome interference.

Another way to demonstrate the importance of ribosome collisions is to directly estimate *t*_*i*_(collisions). Since the total time spent on a codon is given by *r*_*i*_ divided by the ribosomal current, we obtain

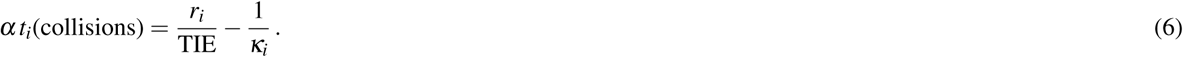

The time spent in traffic on codon *i* is then larger than zero if *r*_*i*_*/*TIE > 1*/κ*_*i*_ and equal to zero only if there is no ribosome interference. In the second row of Figure 7 we show that many of the codons analysed deviate from the bisect. This is another quantitative evidence that, according to experimental data, it is not that rare to observe ribosomes queuing *in vivo*.

### 3.6 Initiation and elongation interdependence

After observing that translation elongation efficiency is generally close to its optimum value of 1, we now look for spatial distribution of the TEE_*i*_ along the transcripts. To this end we compute a metagene TEE profile by aligning the genes at their START codon and computing the distributions of the TEE_*i*_ at each position *i*. We then take the median of the distribution on each codon. The results are plotted in Figure 8(a). This genome-averaged profile confirms our earlier observation that TEE is close to 1. However, we also observe that the first ∼ 10 sites have a larger elongation efficiency. A large value of TEE around the START codon helps to clear this region from queueing ribosomes and thus facilitates ribosome recruitment (see also Relationship between TIE and TEE in Supplementary Information). This result is consistent with a large value of TIE previously observed in Figure 5, and it confirms the importance of the beginning of the coding sequence in controlling translation.

**Figure 8.**
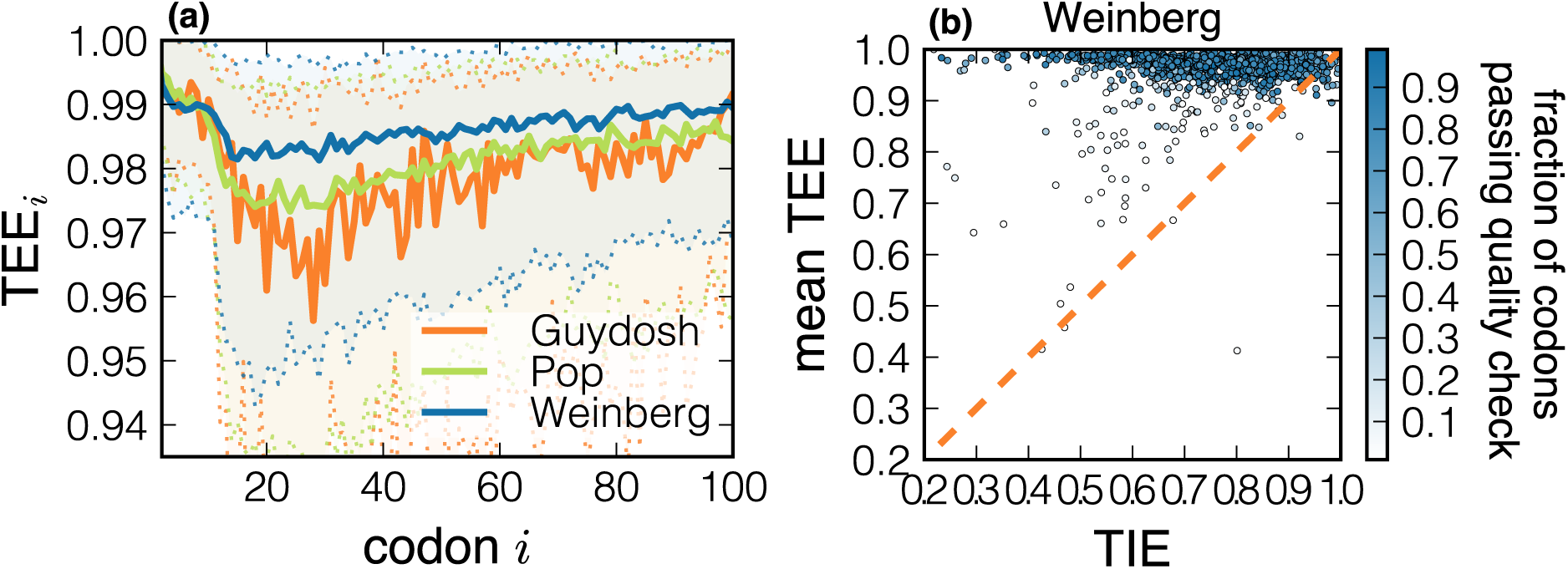
(a) Translation Elongation Efficiency metagene profile. The full line is the median TEE profile for three different datasets included in our study. Dotted lines delimiting the shadow area correspond to the first and third quartiles of the distribution. In panel (b) we show the scatter plot of the TIE and mean TEE of each gene in the Weinberg dataset. The orange dashed line represents mean TEE = TIE. The analysis of the other dataset can be found in Figure S8. Points are colored according to the fraction of codons passing the quality check.

Our results seem to suggest that TIE and TEE should be strongly related. On the one hand, if translation initiation is efficient but elongation is inefficient, ribosome interference would dominate and ribosomal resources would be wasted. On the other hand, effective elongation and weak initiation would still finely tune the overall protein production without harming cellular fitness. Following these considerations, from the evolutionary point of view there should exist a constraint between the relative weights of initiation and elongation, and a situation with strong initiation and weak elongation should be avoided.

We can roughly evaluate the overall elongation efficiency as the mean of the TEE_*i*_ profile of each gene, and thus associate a couple of values (TIE, mean TEE) to each gene analysed. In Figure 8(b) we observe that the constraint TIE *<* mean TEE is satisfied for most of the genes analysed (only a few exceed the TIE = mean TEE dashed line, and for very initiation efficient genes); thus the data analysed are consistent with the hypothesis explained above. We also notice that transcript with inefficient initiation might also present a less optimised elongation, suggesting that initiation and elongation are interdependent.

## 4 Discussion

In this work we introduce NEAR, a Non-Equilibrium Analysis of Ribo-seq, which is based on a well-studied model borrowed from statistical physics. The model tracks individual ribosomes engaged in translation and predicts their spatial distribution on the mRNA and the rate of protein synthesis using initiation, elongation and termination rates as input parameters. Here we do the opposite—we develop a method that infers elongation-to-initiation ratios at codon resolution directly from ribosome profiling data.

We first emphasise that Ribo-seq profiles, being an averaged snapshot of the translatome, do not contain information on the absolute timescales of the process, and that thus it is possible to estimate relative rates only. These rates uniquely determine the density profile and allow us to evaluate the extent of ribosome traffic along the transcript and show a possible interplay between initiation and elongation. To this end we introduce new measures of translation efficiency that we named translation initiation and elongation efficiencies (TIE and TEE, respectively). Importantly, both TIE and TEE are dimensionless scores taking values between 0 and 1, which allows us to compare ribosome traffic between different genes.

Translation initiation efficiency (TIE) is defined as the probability that a ribosome attempting to initiate translation is not obstructed by another ribosome on the coding sequence. The distribution of TIE for the three datasets that we used in this study show that ribosomes can easily access most transcripts, with the median value of 0.8 for the probability to find the initiation region unobstructed (Figure 6). Yet, we find genes with low TIE suggesting that the first codons can exert control over protein synthesis through ribosome traffic interfering with translation initiation. These results are in line with recent experimental evidence on ribosome stalling and traffic in the initiation region^17, 46, 47^.

Similarly, translation elongation efficiency (TEE_*i*_) is defined as the probability that a ribosome at codon position *i* is not blocked by another ribosome downstream of *i*. The distribution of TEE_*i*_ across all transcripts shows that TEE_*i*_ is generally close to 1 suggesting that ribosome interference is negligible for most codons (Figure 6(e)). However, when looking at the individual gene TEE profiles, we observe that it is not so rare to find the probability of ribosome interference as high as 50% (Figures 6(a)-(d)). In accordance with these results, we find more evidence of ribosome interference (lower TEE) in genes with higher ribosome density (Figure 7). We also compute the average time *t*_*i*_(collisions) that each ribosome spends on a codon due to the blockage of downstream ribosomes. If no traffic is present then *t*_*i*_(collisions) = 0. Instead, we observe many codons for which *t*_*i*_(collisions) > 0 (Figure 7).

The fact that the value of TEE at each codon must fall between 0 and 1 allows us to agglomerate all values of TEE into a “metagene” profile (Figure 8). Interestingly, the median TEE shows slightly higher values at the first 10 codons, suggesting that queuing is avoided in order to allow for efficient ribosome recruitment at the start codon. This result is consistent with a recent study in which replacing the first 8 codons with their slower synonymous variants significantly reduced protein expression without affecting mRNA levels^46^. Furthermore, the first codons have been recognised as critical in determining protein synthesis both theoretically^12, 13, 48^ and experimentally^47, 49–51^. Beyond the first 10 codons, the metagene profile of TEE further reveals a small but noticeable drop between codons 10 and 20, followed by a slow increase between codons 20 and 100. These results are consistent with the “ramp hypothesis” proposing that rare codons are more frequent at the beginning of genes in order to avoid ribosome traffic further along the transcript^52^.

All together, our results indicate that translation initiation is slow compared to elongation (all *κ*_*i*_ = *k*_*i*_*/α <* 1) and the coding sequence interfering with initiation is cleared efficiently (median TIE at 0.8). We also find that translation elongation is largely optimised to avoid traffic (median TEE at 0.99), although one can locally observe high levels of ribosome interference. Interestingly, despite variations in translation initiation efficiency between genes (Figure 5), elongation remains consistently more efficient than initiation (mean TEE > TIE, Figure 8). It is possible that the relative role of elongation and initiation is under evolutionary pressure to allow for an efficient ribosome recruitment and to avoid ribosome interference for efficient transition from initiation to elongation^52^.

Perhaps the most surprising result of our study is the variability of the inferred elongation-to-initiation ratios *κ*_*i*_. We can affirm that there is a correlation between common indices of codon optimality, such as the local tAI, and the estimated elongation-to-initiation ratios (see Figure S6). However, the large variability of the estimated rates of each individual codon type implies that using those indices for protein synthesis optimisation or other synthetic applications will probably not lead to the expected results. Instead, our findings indicate that codon context in the sequence is as relevant as the particular codon used, and further studies should focus on the discovery of mechanisms giving rise to the codon context dependence. For instance, mRNA secondary structures might be relevant, particularly around the initiation region^49–51, 53^ or the amino-acid charge at the beginning of the coding sequence^42^.

Our method has detected many codons at which the model is incompatible with the ribosome profiling data, particularly for genes for which we estimated high level of ribosome interference (see for instance Figure S7). One possibility is that our results are affected by known biases during the bioinformatic analysis^54^. Another source of inconsistency between the model and the data is possibly hidden in the nature of the ribosome profiling technique. Queuing ribosomes generate large footprints^14^ that are usually discarded in the experimental pipeline. Intuitively, one would expect that ribosome profiling discarding large footprints would be insensitive to ribosome interference. However, we note that the model is able to capture correlations between ribosomes that are not immediately adjacent to each other. A recent theoretical study by Scott and Szavits-Nossan (2019)^48^ showed that a slow codon affects ribosome density over multiple codons, although the effect subsides with the distance from the slow codon. Indeed, NEAR finds evidence of local jamming despite the experimental bias that discards jammed ribosomes. We remark that the high TIE and TEE values at the first 10 codons could also be attributed to the nature of Ribo-seq that exclude disome footprints; a recent study by Diament *et al*.^17^ in fact showed that the largest concentration of disomes in *S. cerevisiae* is at the first 10 codons.

Finally, we note that some transcripts show a significant number of rejected codons whose estimated *κ*_*i*_ cannot be considered reliable (see Figure S1). In those cases the best estimate we have for *κ*_*i*_ is the mean-field approximation that neglects correlations between closely spaced ribosomes. Consequently, TIE and TEE may become less reliable, too. Interestingly, transcripts with many rejected codons generally display low values of TIE and mean TEE (Figure 8(b)). There seems to be a connection between how well the TASEP fits the ribosome profiling data and the extent of ribosome traffic that needs further investigation.

To summarise, we have developed a model-based method for inferring codon-specific elongation rates (relative to the initiation rate) from ribosome profiling data. In addition, we have proposed new measures of translation initiation and elongation efficiencies that quantify the extent of ribosome traffic *in vivo* and can be used to compare different genes and experimental conditions. We believe these new scores will complement the standard indices of translation efficiency and will contribute to the understanding of this complex biological process.

Despite the tremendous importance and potential of ribosome profiling, our work emphasises its limitations when deciphering translation dynamics such as the lack of quantification in physical units and the lack of absolute time scales. These challenges have been recognised and steps have been made recently to combine Ribo-seq with other methods for absolute quantification such as RNA-seq with spike-ins^1^ and pulsed stable isotope labelling of amino acids^18^. Future developments of NEAR will include these data to obtain a more detailed view on translation dynamics. Another key question that quantitative studies using ribosome profiling should address in the future is the role of density normalisation in order to better compare the outcome of different genes and of different organisms.

## Acknowledgements

We would like to thank Guillaume Cambray and Edward Wallace for useful discussions. JSN was supported by the Leverhulme Trust Early Career Fellowship under grant number ECF-2016-768. This project has been initially supported by the Défi InPhyNiTi (exploratory project funded by the CNRS).

## Author Contributions

LC and JSN conceptualised the study, developed the methodology and wrote the manuscript. JSN performed the stochastic simulations and LC analysed the data.

## Supplementary Information

### A Analytic results in the steady state as a basis of NEAR

The model has been defined in the Materials and Methods section of the main text. Here we explain how to compute the ribosomal current *J* and the local ribosome density *ρ*_*i*_ using two approximate approaches: the mean-field approximation developed in MacDonald *et al*.^11, 31^ and Shaw *et al*.^55^, and initiation-limited approximation developed in Szavits-Nossan et *al*^12, 27^.

The theory summarised in this Supplementary Data constitutes the basis of NEAR, whose procedure is given in the section Non-Equilibrium Analysis of Ribo-seq (NEAR) of the main text.

#### A.1 Ribosome density and current in the mean-field approximation

In the mean-field approximation, correlations between ribosomes are ignored, which leads to the following system of equations for *J* and *ρ*_*i*_,

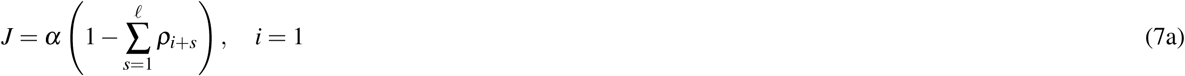

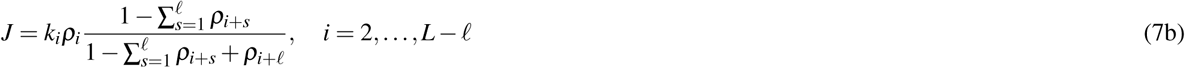

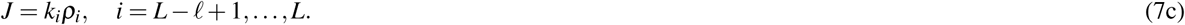

It is straightforward to invert these equations to get the ratio *k*_*i*_/α as a function of the local densities *ρ*_2_; : : : ; *ρ*_L_:

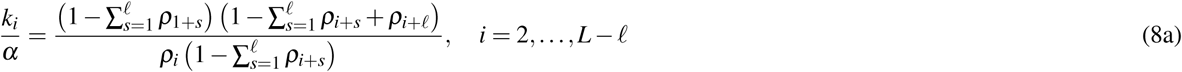

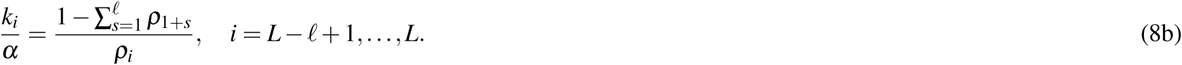

We will use these expressions as a starting point for the nonlinear least-squares minimisation procedure described in Section A, which is the core of NEAR.

#### A.2 Ribosome density and current in the initiation-limited approximation

Recently we introduced a power-series method for computing *J, ρ*_*i*_ and *ρ* by expanding steady-state probability *P*^∗^(*C*) in the initiation rate *α*,

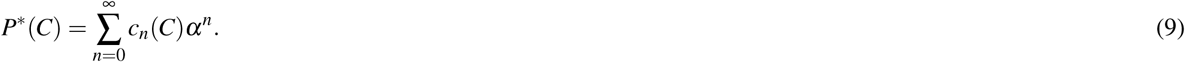

The method then delineates how to compute the coefficients *c*_*n*_(*C*) recursively starting from *n* = 0. Due to its recursive nature, the power-series method is much faster than a stochastic simulation. This is absolutely crucial in order to infer {*k*_*i*_*/α*} from {*ρ*_*i*_}, which is the main part of NEAR.

It turns out that most coefficients *c*_*n*_(*C*) for small *n* are in fact equal to zero, which considerably simplifies the calculation of *P*(*C*). Specifically, if *N*(*C*) denotes the number of ribosomes in a configuration *C* then

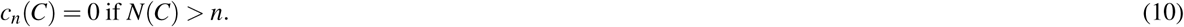

This non-trivial result follows from a graph-theoretical interpretation of Markov chains applied to the TASEP, see Szavits-Nossan et *al*. (2018a)^12^ and Szavits-Nossan et *al*. (2018a)^27^ for more details. Another useful relation is

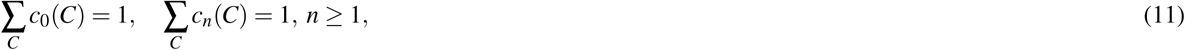

which follows from the fact that the sum of *P*(*C*) over all *C* is equal to 1.

If initiation is slow we can keep the first *K* terms in the series (9) and ignore all the rest, which we call the initiation-limited approximation (ILA),

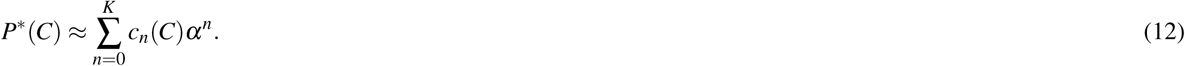

In this work we compute coefficients up to and including the third order (*K* = 3), which we present in detail below. According to Eq. (10), for *K* = 3 we only need to consider configurations with at most three ribosomes on the lattice. These are *C* = ∅ (no ribosomes), *C* = *A*_*i*_ (one ribosome at codon *i*), *C* = *A*_*i*_*A*_*j*_ (two ribosomes at codons *i* and *j*) and *C* = *A*_*i*_*A*_*j*_*A*_*k*_ (three ribosomes at codons *i, j* and *k*).

For *n* = 0 (zeroth order), *c*_0_(*C*) = 1 if the lattice is empty (*C* = ∅) and is equal to 0 otherwise, which leads to *P*(*C*) = 1 if we ignore higher-order terms. This is equivalent of saying that if translation initiation is not allowed (*α* = 0), then the transcript will become completely empty with probability 1.

For *n* = 1 (first order), *c*_1_(*C*) ≠ 0 only if *C* contains at most one ribosome. The corresponding coefficients *c*_1_(∅) and *c*_1_(*A*_*i*_) are equal to

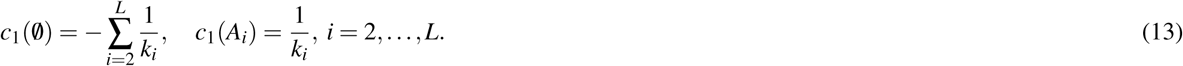

These coefficients yield the following expressions for *J, ρ*_*i*_ and *ρ* in the first-order approximation

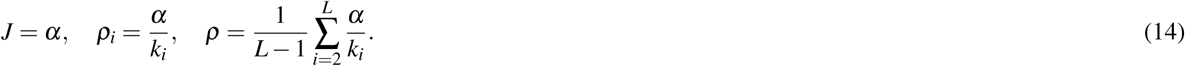

We note that the relationship *ρ*_*i*_ ≈ *α/k*_*i*_ between ribosome densities and elongation rates that we derived above is often used in the interpretation of ribosome profiling data. Because the first order considers configurations with at most one ribosome per lattice, this result neglects the interference between ribosomes and is thus valid only for very small initiation rates.

For *n* = 2 (second order), *c*_2_(*C*) ≠ 0 only if *C* contains at most two ribosomes. The equations for *c*_2_(*C*) are more involved than for *c*_1_(*C*) and must be solved numerically. Before we write down the equations, we introduce Kronecker delta function *δ*_*i j*_ and unit step function *θ* (*i*) defined as

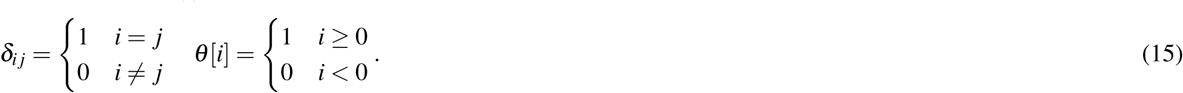

The equations for two-particle coefficients *c*_2_(*A*_*i*_*A*_*j*_) then read

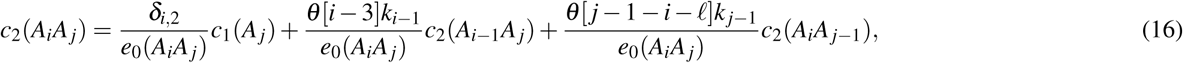

where *e*_0_(*A*_*i*_*A*_*j*_) = *θ*_*j*−*ℓ*−*i*−1_*k*_*i*_ + *k* _*j*_. These equations can be solved recursively starting from *i* = 2 and iterating over *j* from *j* = *ℓ* + 2 to *j* = *L* with *i* held fixed. Then *i* = 3 is held fixed and the iteration goes over *j* from *j* = *ℓ* + 3 to *L* and so on until *i* = *L* − *ℓ* and *j* = *L*. Once all *c*_2_(*A*_*i*_*A*_*j*_) are found, the equations for one-particle coefficients *c*_2_(*A*_*i*_) are solved recursively from *i* = 2 to *i* = *L*,

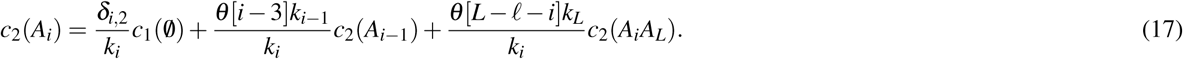

Finally, once all *c*_2_(*A*_*i*_*A*_*j*_) and *c*_2_(*A*_*i*_) are computed, *c*_2_(0/) follows from Eq. (11),

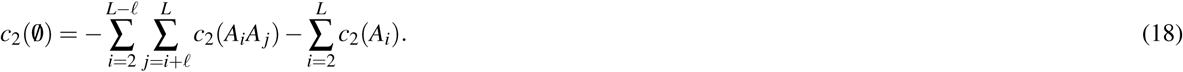

Using these coefficients we can compute *J* and *ρ*_*i*_ in the second-order approximation according to

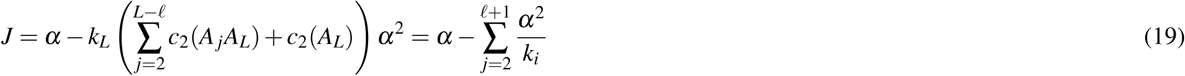

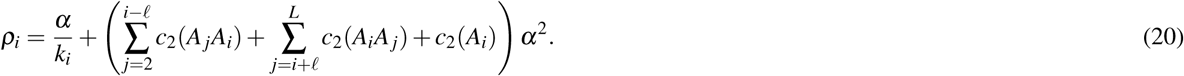

For *n* = 3 (third order), *c*_3_(*C*) ≠ 0 only if *C* contains at most three ribosomes. The equations for three-particle coefficients *c*_2_(*A*_*i*_*A*_*j*_*A*_*m*_) read

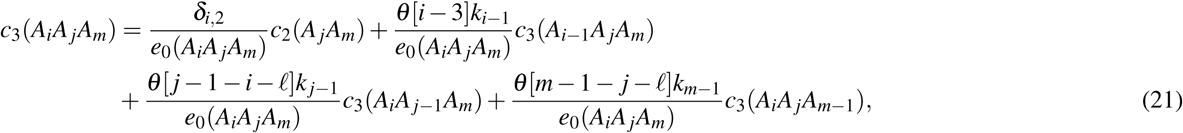

where *e*_0_(*A*_*i*_*A*_*j*_*A*_*m*_) is given by

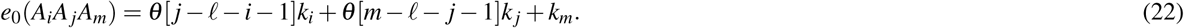

These equations are solved recursively starting from *i* = 2 and *j* = *ℓ* + 2 fixed and iterating over *m* from 2*ℓ* + 2 to *L*. Then *j* is increased to *ℓ* + 3 and the iteration over *m* is repeated from 2*ℓ* + 3 to *L*. The procedure of increasing *i* and *j* and iterating over *m* is repeated so on until *i* = *L*− 2*ℓ, j* = *L*−*ℓ* and *m* = *L*.

Once all *c*_3_(*A*_*i*_*A*_*j*_*A*_*m*_) are found, the equations for two-particle coefficients *c*_3_(*A*_*i*_*A*_*j*_) are solved recursively starting from *i* = 2 and *j* = *ℓ* + 2,

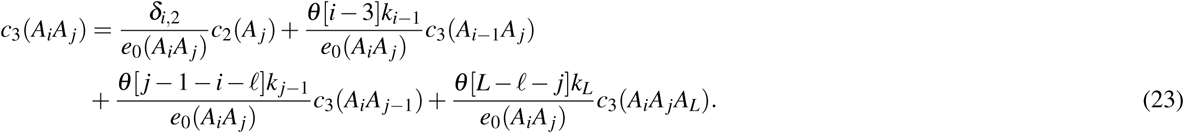

The equations for one-particle coefficients *c*_3_(*A*_*i*_) are then solved recursively from *i* = 2 to *i* = *L*,

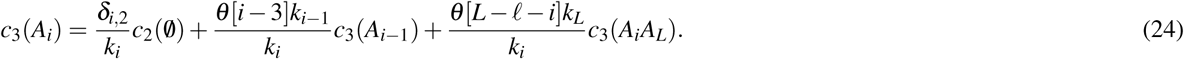

Finally, the coefficient *c*_3_(∅) is given by

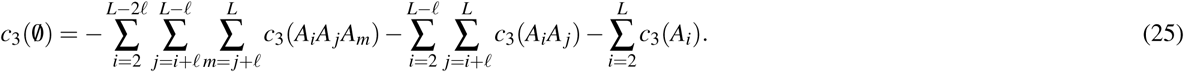

Once all coefficients for all orders up to and including the third order are computed, *J* and *ρ*_*i*_ can be computed according to the following expressions

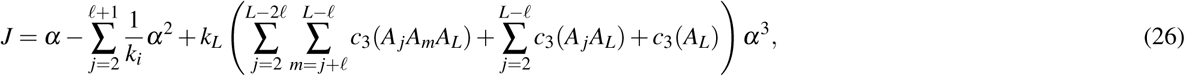

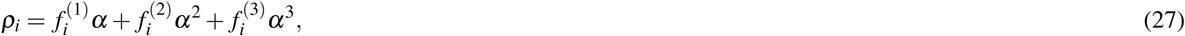

where the coefficients 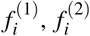 a nd 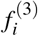 depend only on *k* _2_, …, *k*_*L*_ and are given by

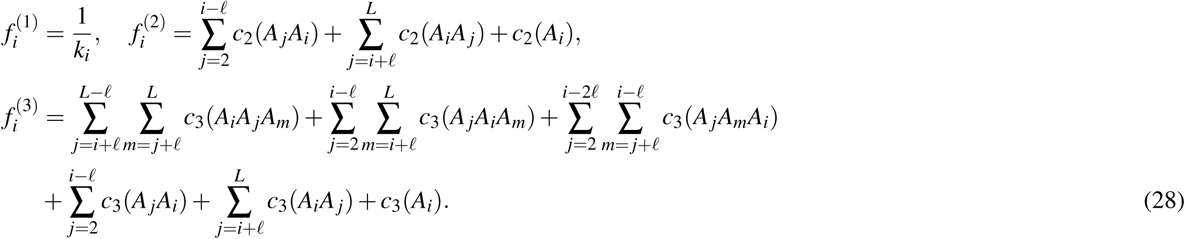

From the stationary master equation in the Material and Methods section of the main text, it follows that *P*(*C*) and thus *ρ*_*i*_ are functions of {*κ*_*i*_} = {*k*_*i*_*/α*} only. Using this fact we can absorb *α*^*n*^ into *f* ^(*n*)^(*C*) and rewrite Eq. (27) as

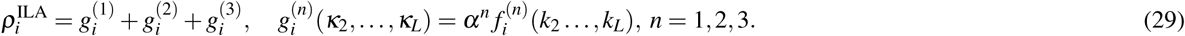

### B Details of the NEAR method

The method of inferring elongation-to-initiation ratios from ribosome profiling data is summarised in the Section Non-equilibrium Analysis of Ribo-seq (NEAR) of the main text. Here we provide further details of the stochastic simulations, absolute normalisation of Ribo-seq data and non-linear optimisation for computing elongation-to-initiation ratios {*κ*_*i*_}.

#### B.1 Stochastic simulations

All Monte Carlo simulations were performed using the Gillespie algorithm. In the first part of the simulation we checked the total density *ρ* every 100 · *L* updates until the percentage error between two values of the total density *ρ* was less than 0.1%. After that we ran the simulation for further *M* = 10^4^ · *L* updates during which we computed the time average of *ρ*_*i*_ defined as

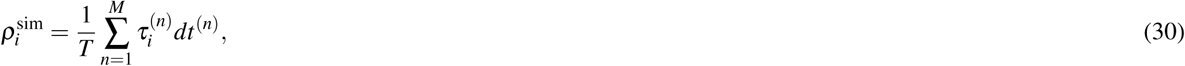

where 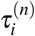 is the value of *τ*_*i*_ (1 if codon *i* is occupied by the ribosome’s A-site and 0 otherwise) just before the *n*-th update in the simulation, *dt*^(*n*)^ is the time interval between the (*n* − 1)-th and the *n*-th iteration of the Gillespie algorithm, and 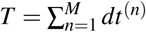 is the total time.

### B.2 Normalisation of Ribo-seq data

Genes from the three datasets analysed were selected to have at least 10 “reads” per codon on average. For each gene we computed the value of local experimental ribosome density *r*_*i*_ at codons *i* = 2, …, *L* according to

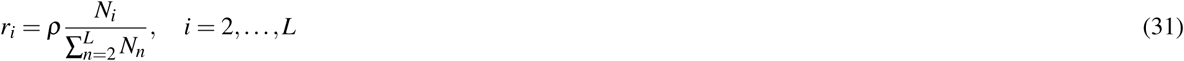

where *N*_*i*_ is the number of reads at codon *i* obtained from ribosome profiling experiments, *ρ* is the total ribosome density and *L* is the number of codons for that particular gene. Most genes in our analysis had codons with zero reads, which implies an infinite value of the elongation rate. In order to mitigate this problem we replaced *N*_*i*_ = 0 by *N*_*i*_ = 1 and calculated *r*_*i*_ according to Eq. (31). The normalisation procedure failed for only three genes across all three datasets, because the normalised ribosome density *r*_*i*_ was larger than 1 at some codons.

### B.3 Non-linear optimisation for computing elongation-to-initiation ratios

We looked for *κ*_*i*_ = *k*_*i*_*/α* such that the theoretical ribosome density 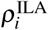 predicted from the initiation-limited approximation in Eq. (27) matched the experimental density *r*_*i*_,

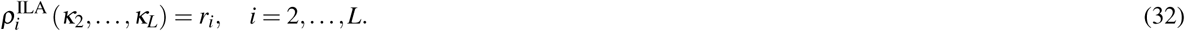

The procedure of finding *κ*_*i*_ consisted of four steps:

1. First we estimated the values of 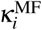 in the mean-field approximation according to Eq. (8).
2. Next we computed local ribosome densities 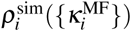 by running a stochastic simulation of the TASEP using the Gillespie algorithm in which the elongation rate *k*_*i*_ at codon *i* was set to 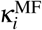 estimated in the mean-field approximation (i.e. setting *α* = 1 in the simulation without loss of generality).
3. We then compared 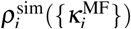 to *r*_*i*_ for each *i* = 2, …, *L* and identified codons *x*(1),…, *x*(*N*) for which the percentage error between 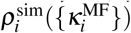 and *r* was larger than 5%.
4. Finally, we solved a nonlinear least squares optimisation problem using 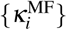 as a starting point, which consisted of finding *κ*_*x*(1)_, …, *κ*_*x*(*N*)_ for which the sum of squares

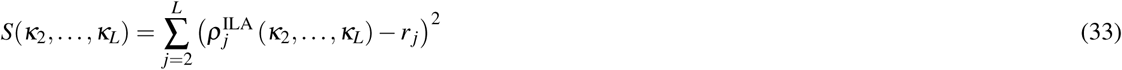

was minimal, where 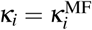 for codons *i* ≠ *x*(1),…, *x*(*N*) that were not selected for optimisation and 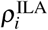 is given in Eq. (29).

The nonlinear optimisation was preformed using NLopt library from Ref.^56^. We used a local derivative-free algorithm called BOBYQA which was developed in Ref.^57^. In order to prevent unrealistic values of *κ*_*i*_ and to speed up the optimisation procedure, we restricted the search to 10^−3^ ≤ *κ*_*x*(1)_, …, *κ*_*x*(*N*)_ ≤ 10^6^. The optimisation search was preformed until one of the following two stopping criteria was met: (1) the fractional error Δ*S/S* between two iterations was less than 10^−8^ and (2) the total run time exceeded 60 minutes. These numbers were chosen due to time constraints when analysing many genes; a better accuracy may be achieved for individual genes by amending the stopping criteria. After the optimisation procedure finished we recomputed local densities 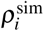 by running a stochastic simulation of the TASEP with elongation rates *k*_*i*_ = *κ*_*i*_ and compared them to experimental values *r*_*i*_.

### B.4 Quality check of the estimated elongation-to-initiation ratios

In this section we detail the quality check that we carried out for each value of the inferred *κ*_*i*_ = *k*_*i*_*/α*. For each gene for which the normalisation of Ribo-seq data was successful we preformed the following seven checks:

#### 1. Improvement over the initial (mean-field) prediction

We check whether the optimisation procedure has improved the agreement with respect to the initial (mean-field) prediction. For the mean-field prediction we compute the sum of squares *S*_MF_

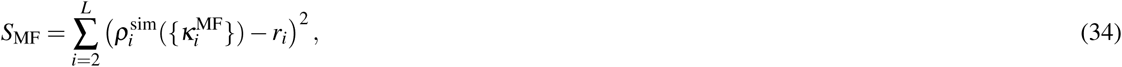

where 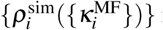 is the simulated density profile computed with the mean-field rates 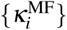 and *r*_*i*_ is the experimental density profile. The value of *S*_MF_ is then compared to *S*_opt_ obtained from

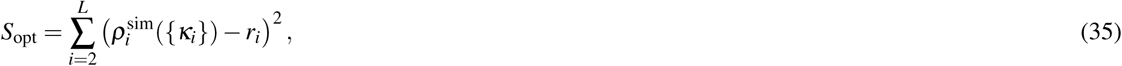

where 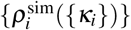 is the simulated density profile computed using the inferred elongation-to-initiation ratios {*κ*_*i*_}. If *S*_opt_ *< S*_MF_ then the TIE is taken from the simulations of the optimised system, otherwise from the MF simulations.

#### 2. Rate-limiting step in translation

We verify if the optimised elongation-to-initiation ratio *κ*_*i*_ > 1 ∀ *i* (otherwise initiation is not the limiting step and the initiation-limited approximation cannot be used).

#### 3. Applicability of the initiation-limited approximation

We set a tolerance *ε*_ILA_ for the ribosome density 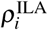 predicted by the initiation-limited approximation in Eq. (27). For each codon we check if 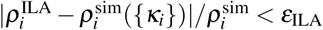. If that is the case the power series approximation holds and the method is reliable. We set *ε*_ILA_ = 0.1.

#### 4. Comparison with Ribo-seq data

If the codon passes the quality checks in points 2 and 3 then we check if the prediction is consistent with the experimental profile {*r*_*i*_} (within a tolerance *ε*_EXP_) by checking if 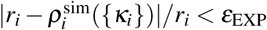, where we set *ε*_EXP_ = 0.05. Codons that pass this check are kept and considered reliable only if *κ*_*i*_ *< κ*_thr_. This last check is to discard values of *κ*_*i*_ that are deemed suspiciously large and are likely unrealistic. Such unreasonably large values of *κ*_*i*_ are due to low number of reads for a particular codon, especially for codons for which we artificially increased the number reads from zero to one in order to avoid infinite elongation rates. Because such small number of reads may be due to experimental errors, we decided to exclude those codons from the final analysis. We used the threshold value *κ*_thr_ = 500 corresponding to an elongation rate *k*= 60/s (assuming *α* = 0.12/s).

#### 5. Problematic codons

If 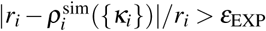 then the codon is excluded from the final analysis and we set *κ*_*i*_ = − 1 (to identify the problematic codon for further analysis). Those codons fall in the most interesting class, in which the experimental data cannot be reproduced using the existing theory. We speculate that those “problematic” codons might also arise because ribosome profiling cannot detect clusters of ribosomes that will instead be predicted by our simulated profile 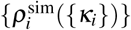.

The fraction of problematic codons for each of the three datasets is presented in Figure S1. A full list of codons entering in this category (in which our method can be applied but experimental data are inconsistent with the density generated by NEAR) is in the Supplementary Files 1,2,3.

#### 6. Falling back to the mean-field prediction

In the case in which the codon does not pass the quality checks in points 2 and 3, the mean-field prediction 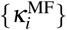 will be considered. If 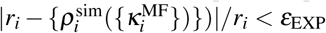, then 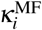 is kept and the mean-field prediction is considered reliable. Otherwise the codon is excluded from the final analysis and we set *κ*_*i*_ = −2.

#### 7. Stop codon

We further check if the elongation-to-initiation ratio *κ*_*L*_ of the stop codon has been kept for the analysis. If *κ*_*L*_ is reliable then the ratio *κ*_*i*_*/κ*_*L*_ = *k*_*i*_*/k*_*L*_ can be computed.

### B.5 Computing TEE profiles

We remind the following relation from the main text: *t*_*i*_(total) = *t*_*i*_(intrinsic) + *t*_*i*_(collision). By definition of TEE_*i*_,

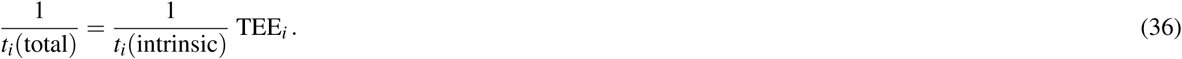

**Figure S1.**
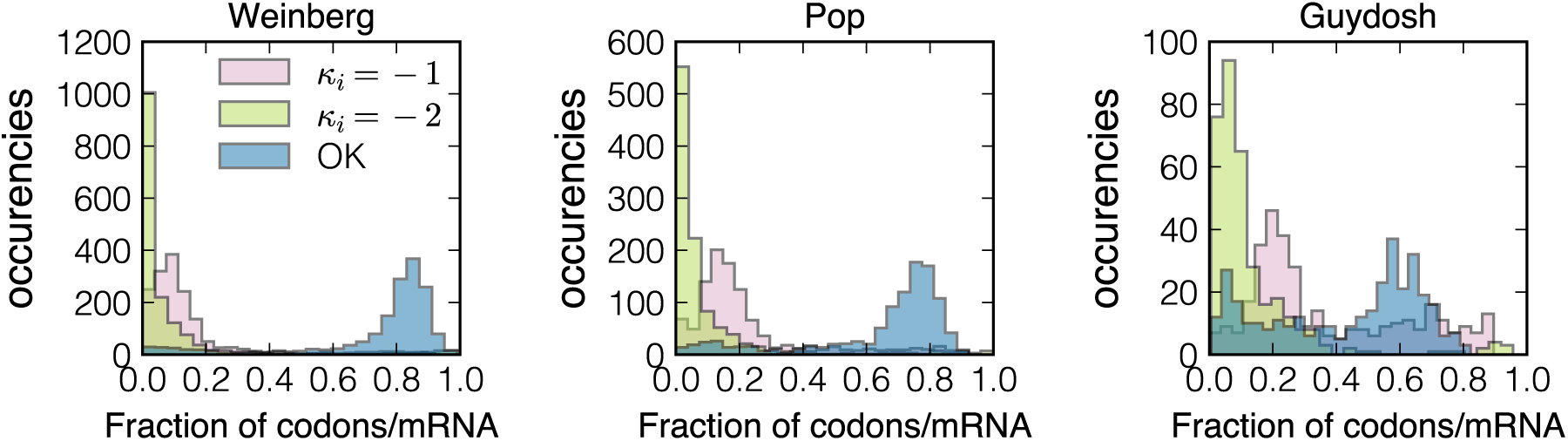
In pink we show the distribution of the fraction of codons per mRNA which are considered problematic (step 5, or *κ*_*i*_ = − 1), while in green we represent the distribution of the fraction of codons per transcript that do not pass step 6 (i.e. for which the initiation-limited approximation is not valid and the mean-field is not considered reliable neither, *κ*_*i*_ = − 2). The equivalent distribution for the fraction of codons passing the quality check is in blue. We remind that the number of analysed genes in the Weinberg, Pop and Guydosh datasets are 1589, 1051 and 345 respectively. The median fraction of codons/mRNA passing the quality check is 0.83, 0.73, 0.51 for these datasets.

The ribosomal current is given by

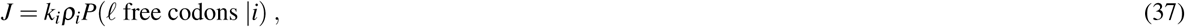

where *P*(*ℓ* free codons |*i*) is the probability that the *i* + 1 … *i* + *ℓ* codons are not occupied (given that a ribosome’s A-site is at site *i*). If we identify the elongation rate *k*_*i*_ with 1*/t*_*i*_(intrinsic) and *J/ρ*_*i*_ with the effective elongation rate 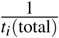, then by comparing Eqs. (36) and (37) we find that the TEE_*i*_ is equivalent to *P*(*ℓ* free codons | *i*). Based on the definition of the TIE=*J/α*, we obtain the TEE profile by computing 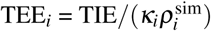 at each codon *i* at which *κ*_*i*_ passed the quality check. Therefore the profile is “broken” when NEAR cannot find a reliable estimate for the elongation-to-initiation ratio *κ*_*i*_. We note that by definition TEE_*i*_ is a probability and as such must take values between 0 and 1. Because each of TIE, *κ*_*i*_ and 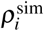 comes with its own statistical error, TEE may occasionally be larger than 1. In those cases we capped the value to 1.

### B.6 Relationship between TIE and TEE

Translation initiation index measures the unoccupancy of the first *ℓ* = 10 codons and is thus related to *κ*_*i*_ and TEE_*i*_ in that region. The TIE is defined as *J/α*, where *J* is the ribosome current i.e. the number of successful translation initiations per unit time. The ribosome current is given by *J* = *αP*(first *ℓ* codons free), where *P*(first *ℓ* codons free) is the probability that the first *ℓ* = 10 codons are free. The latter is given by

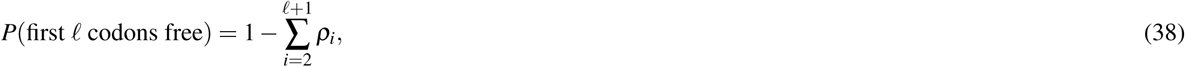

From there we conclude that

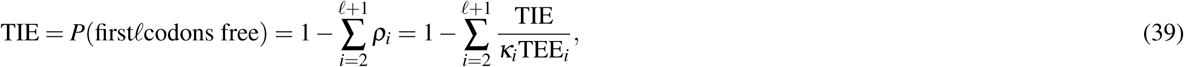

where in the last step we used that *ρ*_*i*_ = TIE*/*(*κ*_*i*_TEE_*i*_). Finally, the relationship between TIE, *κ*_*i*_ and TEE_*i*_ of the first *ℓ* codons is given by

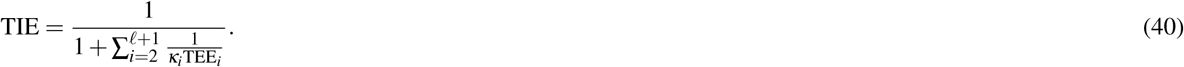

This expression reveals that efficient translation initiation requires both fast elongation (*κ*_*i*_ > 1) and no ribosome queuing (TEE_*i*_ close to 1) within the first *ℓ* = 10 codons.

**Figure S2.**
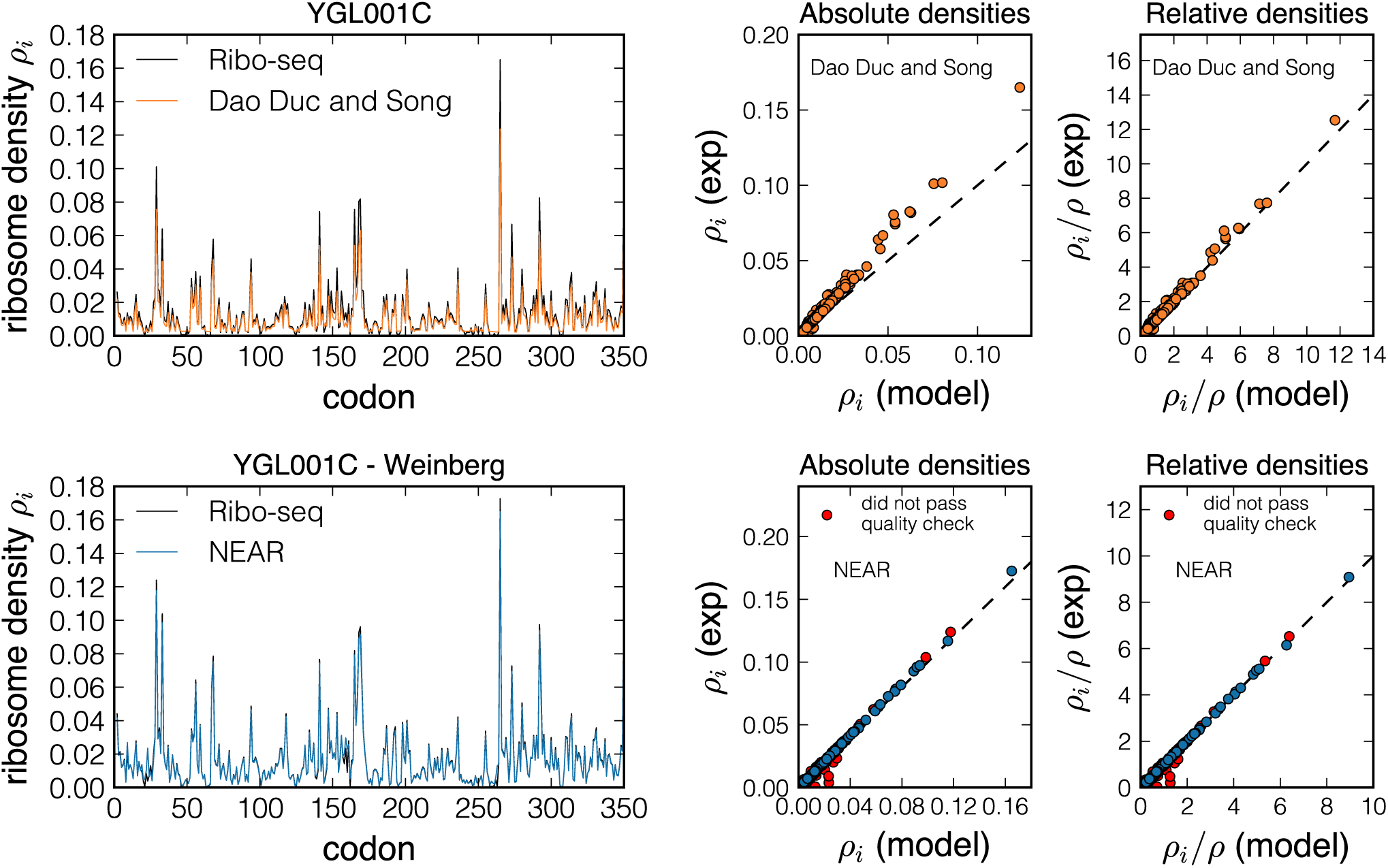
We compare results from Dao Duc and Song^16^ (first row) and NEAR (second row). Although the density profile from simulations with rates obtained by optimising the relative density profile {*ρ*_*i*_*/ρ*} looks comparable with the experimental one (top-left panel), there is a bias between absolute densities (top-middle panel). This follows from the fact that {*ρ*_*i*_*/ρ*} remains unchanged if we multiply all *ρ*_*i*_ by a constant factor, which in turn means different densities *ρ*_*i*_ and therefore different *κ*_*i*_. In the bottom-left panel, ribosome density for the same gene predicted by NEAR (blue line) is compared to the experimental data (black line). NEAR is founded on the optimisation of *absolute* densities, thus both absolute and relative densities match the experimental data (bottom-middle and right panel). In the top row we computed absolute densities as *TE/*(100 × 0.83) × *L* × *R*_*i*_*/R*, where TE is the translation efficiency, *L* is the length of the mRNA in codons, *R*_*i*_ is the number of reads for codon *i* and *R* is the total number of reads of that gene; data from Dao Duc and Song^16^. In the bottom row the experimental densities are the *r*_*i*_ introduced in the main text. We highlight in red (bottom-middle and right panel), the codons that did not pass the quality check of our method.

**Figure S3.**
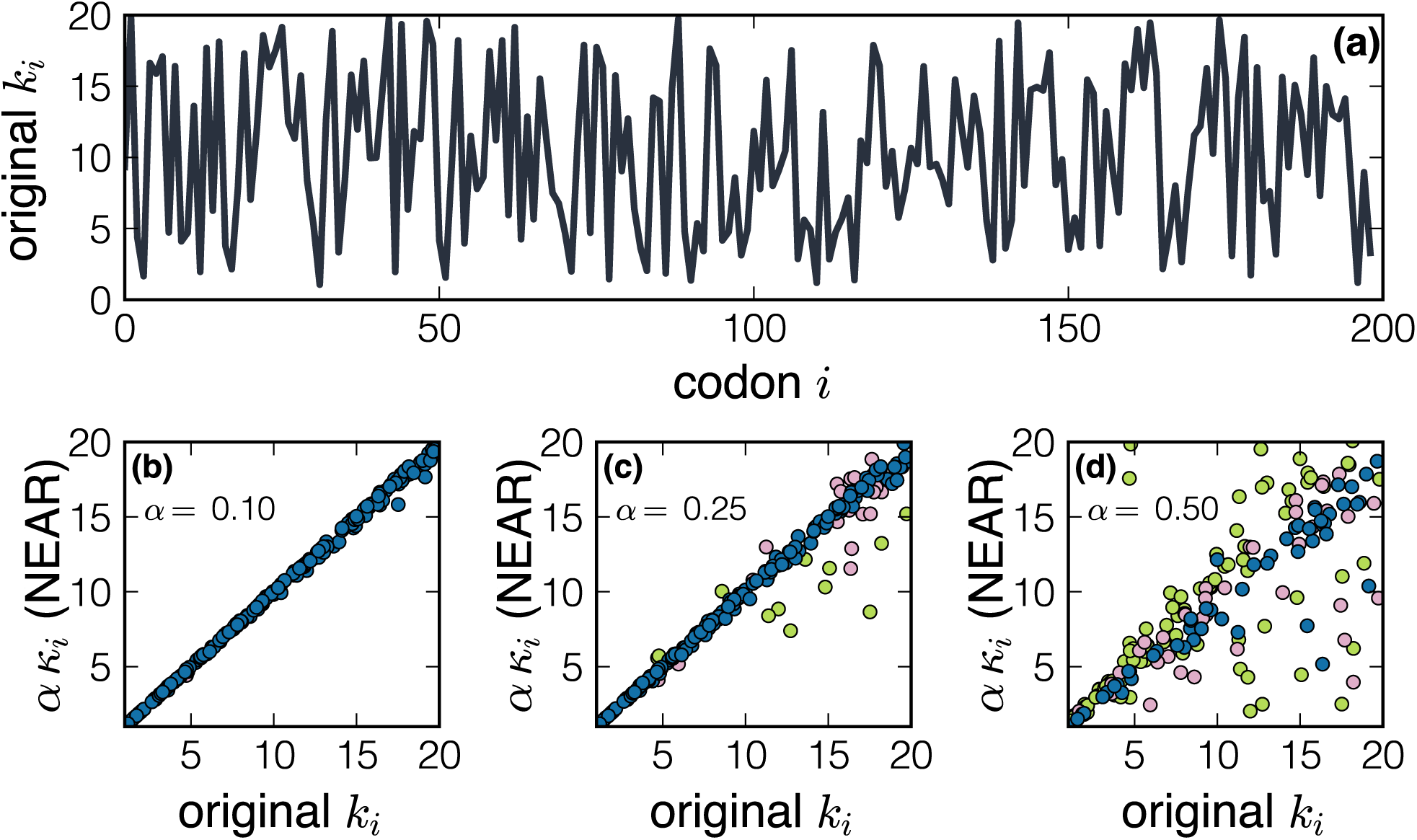
Testing NEAR on a random sequence of rates {*k*_*i*_}. The {*k*_*i*_} profile is shown in panel (a). Panels (b)-(c)-(d) show the scatter plot between the rates *α κ*_*i*_ estimated by NEAR (y-axis) and the original rates {*k*_*i*_} (x-axis) with increasing values of the initiation rate *α*. The points in green are the *κ*_*i*_ values that did not pass the point 4(a) of the quality check as described in the main text (i.e. for which *κ*_*i*_ is fixed to − 2 in the algorithm. Pink points are the ones that did not pass point 4(b), and for which *κ*_*i*_ is arbitrarily fixed to be − 1. Since NEAR is based on a power series approximation that is reliable for small initiation rates, the method is expected to perform badly for large values of *α*. However, even in this regime the quality check is able to exclude codons that will not be considered in the final analysis. We remind that initiation has been estimated to be limiting and the physiological situation is the one presented in panel (b) for most of the genes^37^. For instance in panel (d) one would measure values of *κ*_*i*_ as small as ∼ 2.5, while the values of *κ*_*i*_ estimated from datasets are generally at least on order of magnitude larger (see Figure S4).

**Figure S4.**
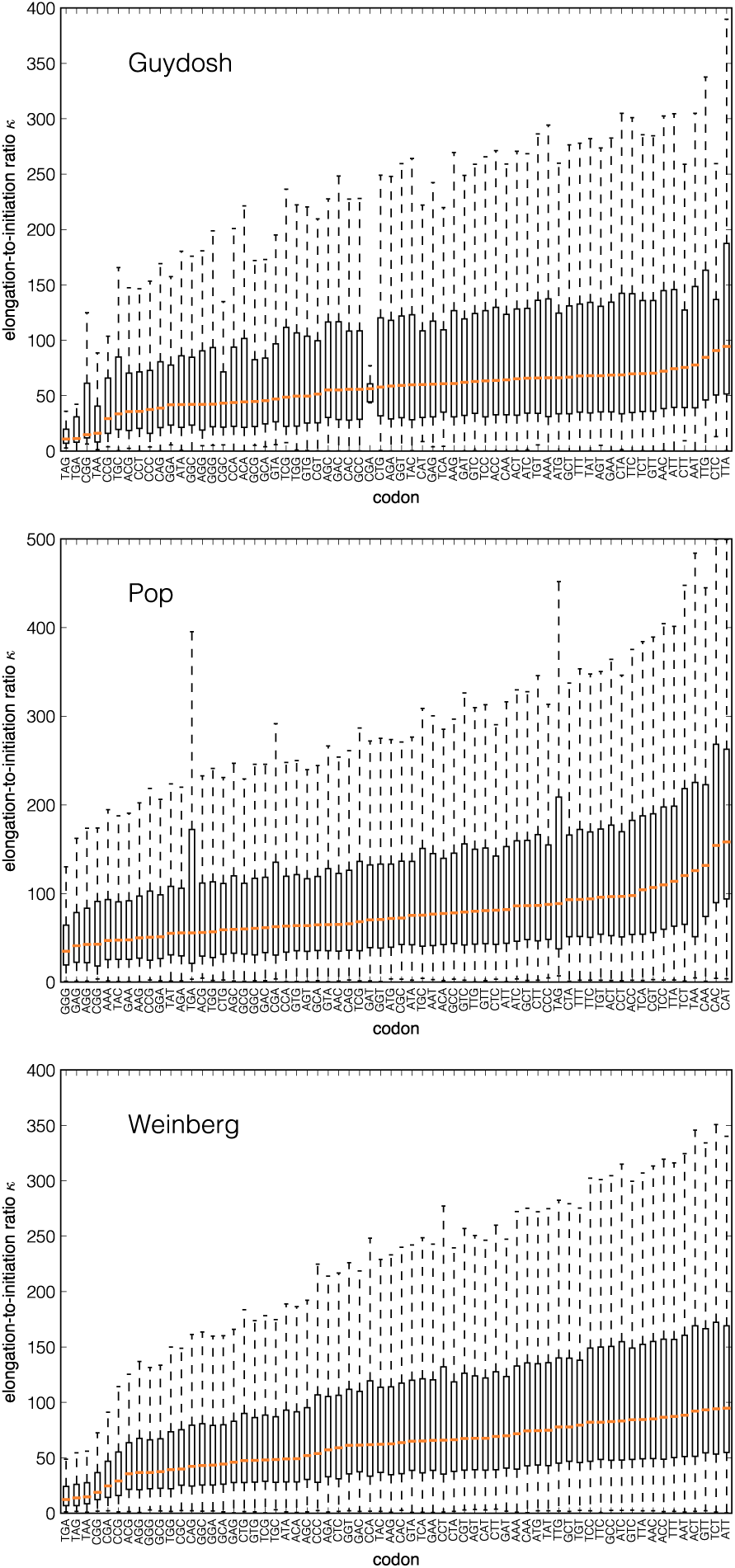
Distributions of elongation-to-initiation ratios *κ* for all codon types. We gather the ratios *κ*_*i*_ = *k*_*i*_*/α* for each codon-type, and plot their distributions. In principle codons from different genes cannot be compared because they have a different initiation rate *α*. However, we notice a small variability in the estimates of STOP codons (TAA, TAG, TAA) for the Guydosh and Weinberg datasets. This is reasonable since STOP codons are supposed to be less context dependent. We also remark that, as opposed to what is generally believed, STOP codons are the slowest, but still ∼ 10 times faster compared to initiation. The boxplots show the quartiles *Q*1 and *Q*3 (Δ*Q* = *Q*3 − *Q*1) of the distribution, the median is the orange horizontal line; the whiskers extend to the most extreme, non-outlier data points (i.e. to the last data point less than *Q*3 + 1.5Δ*Q* and greater than *Q*1 − 1.5Δ*Q*). The Guydosh dataset is the smallest one, and only 10 CGA codons passed the quality check.

**Figure S5.**
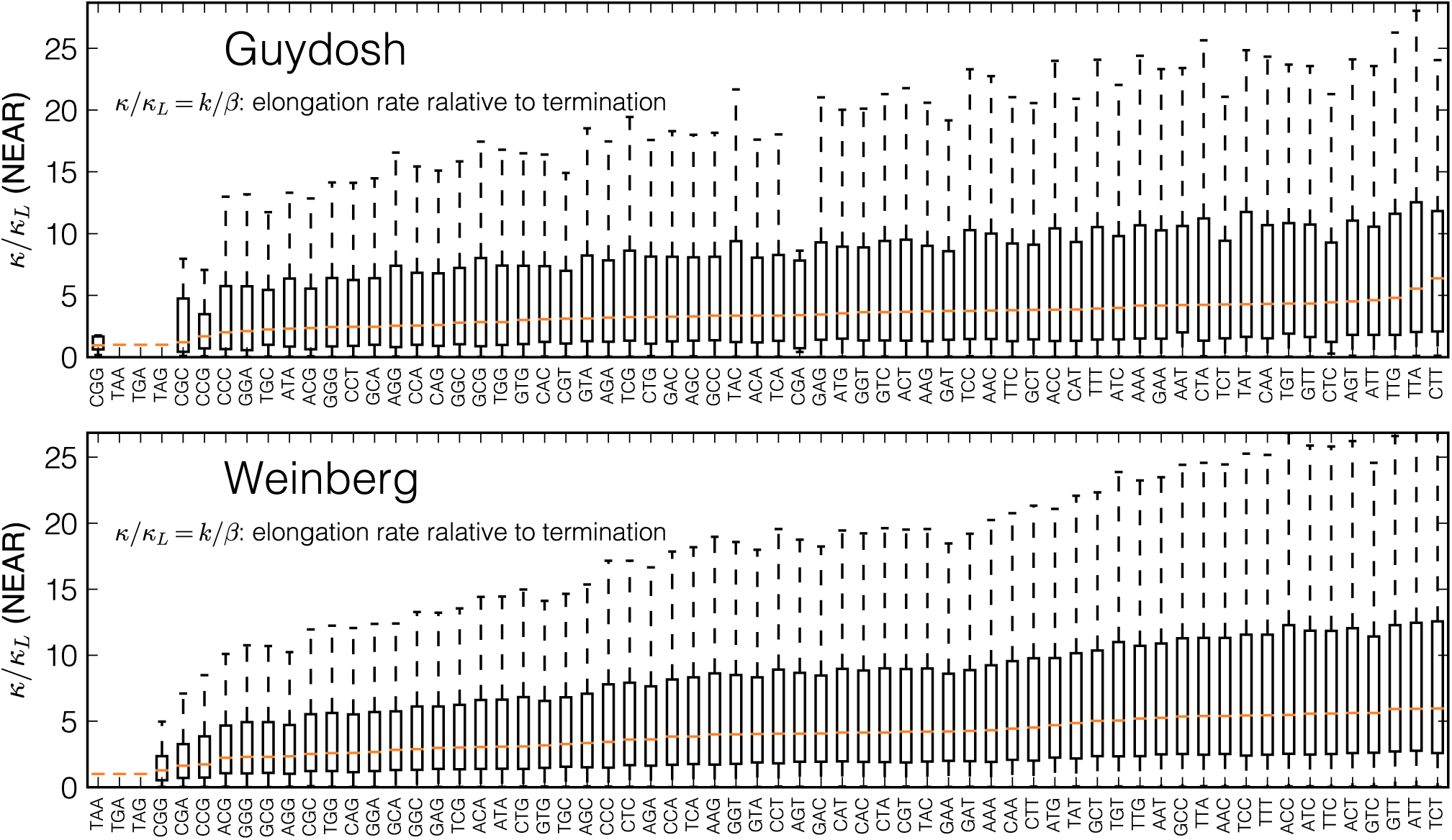
Boxplot of the distribution of the elongation-to-termination ratios *κ/κ*_*L*_ for each codon type for the Guydosh and Weinberg dataset. There is a ≈ 5-fold difference in the median value of *κ/κ*_*L*_ between the slowest and the fastest codon, in accordance with a common view that translation elongation is a non-uniform process. However, in we also find a large variability in the values of *κ/κ*_*L*_ belonging to the same codon type. This variability suggests that the elongation speed of individual codons is only partially determined by their codon type. The boxplots show the quartiles *Q*1 and *Q*3 (Δ*Q* = *Q*3 − *Q*1) of the distribution, the median is the orange horizontal line; the whiskers extend to the most extreme, non-outlier data points (i.e. to the last data point less than *Q*3 + 1.5Δ*Q* and greater than *Q*1 − 1.5Δ*Q*). Only 11 CGG codons of the Guydosh dataset can be used to plot these distributions (both CGG and STOP codon passed the quality check).

**Figure S6.**
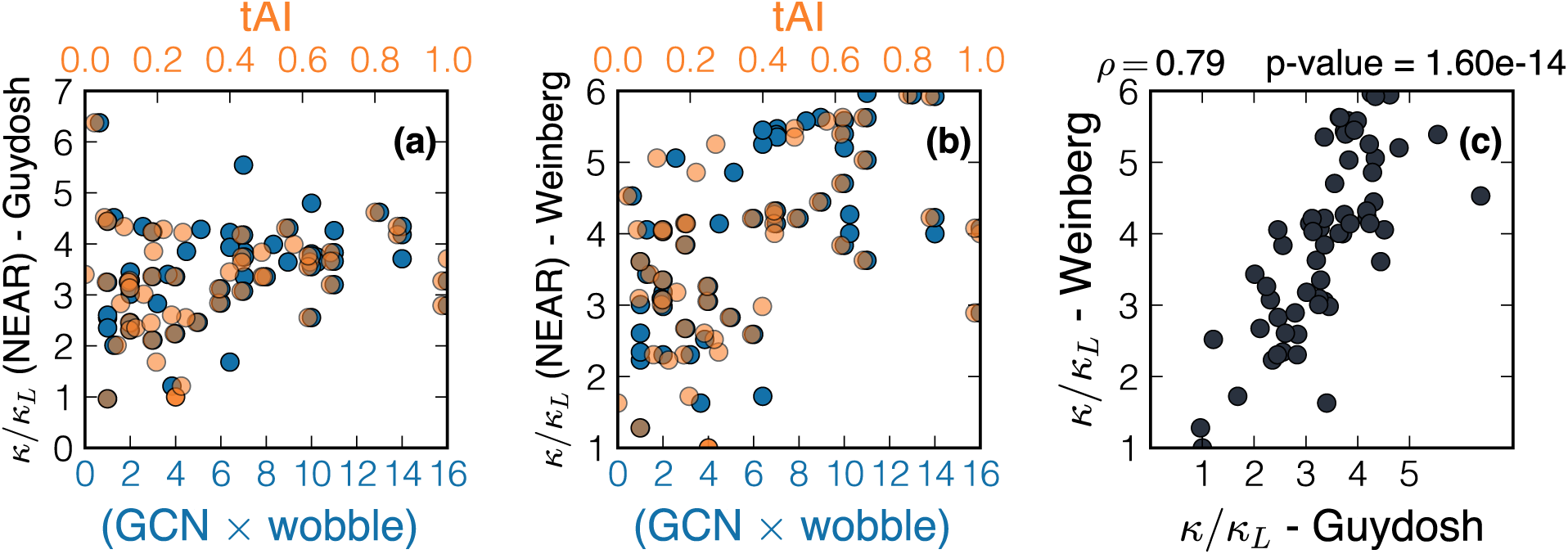
We compare median values of *κ/κ*_*L*_ against two common measures of tRNA availability, a codon-dependent rate of translation based on the tRNA gene copy number (GCN) corrected for the wobble base pairing from Weinberg *et al*.^23^, and the tRNA adaptation index (tAI)^45^ for Guydosh (a) and Weinberg (b) datasets. We find a moderate correlation between the median of the *κ/κ*_*L*_ distributions and both measures of tRNA availability (GCN × wobble: Spearman *ρ* = 0.34, *p* = 0.008 - Guydosh; *ρ* = 0.57, *p* = 1.4 × 10^−6^ - Weinberg; tAI: *ρ* = 0.22, *p* = 0.1 - Guydosh; *ρ* = 0.46, *p* = 3 × 10^−4^); this suggests that the elongation speed of individual codons is only partially determined by their codon type. Panel (c) shows the comparison between the two datasets.

**Figure S7.**
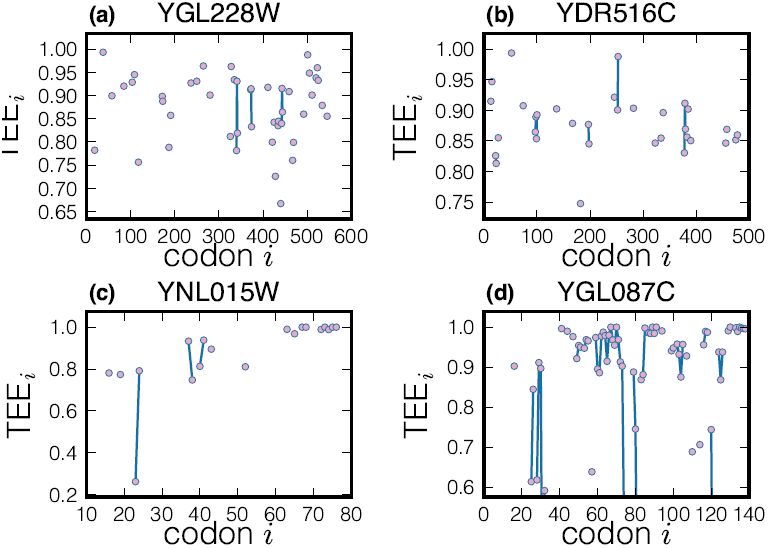
Profiles of TEE for four genes amongst the ones with the smallest average TEE. When the TEE is low, many *κ*_*i*_ do not pass the quality check, as it can be seen from the many points missing in the TEE profiles. This means that the experimental read counts are inconsistent with the model. We speculate that this is due to the bias in the ribosome profiling neglecting clusters of ribosomes. However, NEAR fills the partial information that is enclosed in the experimental profile and finds evidence of traffic (small TEE).

**Figure S8.**
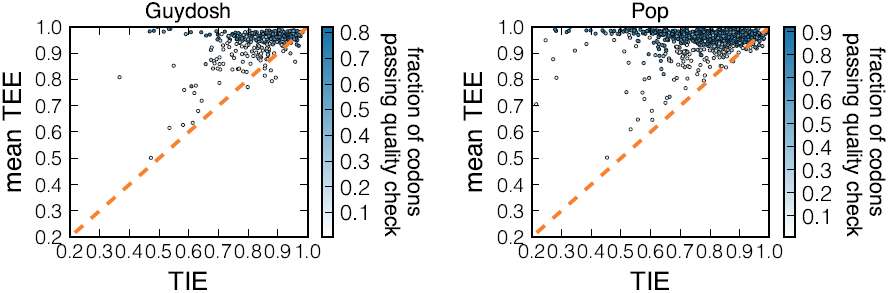
TIE vs mean TEE for the two datasets not shown in the main text.

